# *Acinetobacter* phages use distinct strategies to breach the capsule barrier

**DOI:** 10.1101/2025.07.10.664161

**Authors:** Alexis J McCalla, Forrest C Walker, Fabiana Bisaro, Miguel Rodriguez-Anavitate, Anna Johannesman, Gisela Di Venanzio, Mario F Feldman, Michele LeRoux

**Affiliations:** Department of Molecular Microbiology, Washington University in Saint Louis School of Medicine, Saint Louis, MO 63110, USA

## Abstract

*Acinetobacter baumannii* is an opportunistic pathogen that is a growing threat in hospital settings due to its alarmingly high rates of antibiotic resistance. Alternative therapies are urgently needed to manage the growing burden of untreatable *A. baumannii* infections. Phage therapy is a promising avenue that has already seen some success in isolated compassionate-use cases, including the famous “Patterson case”. *A. baumannii* capsule is highly diverse both in structure and composition, and provides the first immunity barrier against phages. Here, we perform a detailed molecular characterization of three recently isolated, distinct *A. baumannii* phages that breach the capsule via different mechanisms. Like many previously described *A. baumannii* phages, a specific capsule type is necessary and sufficient for StAb1 infection. We found that StAb2 and its relatives adsorb to either a specific capsule type or the conserved outer membrane protein CarO, a porin normally occluded by the capsule. Thus, this phage has a narrow host range amongst capsulated strains, but can broadly infect *A. baumannii* strains lacking capsule. We also show that an unclassified siphophage, StAb3, requires a conserved and uncharacterized glycan, likely containing ManNAc, that enables StAb3 to infect a broad range of *A. baumannii* strains without depolymerizing the capsule. We demonstrate how rationally combining phages with distinct capsule interactions reduces the rapid emergence of phage escape mutants, with potential applications for more effective phage therapy.

## Introduction

The opportunistic pathogen *Acinetobacter baumannii* causes severe, frequently untreatable nosocomial infections that are increasingly challenging to manage due to rising levels of multi-drug resistance (MDR)[1]. Accordingly, both the World Health Organization and the United States Centers for Disease Control and Prevention have determined that developing new treatments for MDR *A. baumannii* is an urgent priority[2,3]. The use of bacteriophages (phages) to treat bacterial infections has reemerged as a promising alternative approach for combating MDR pathogens[4–6]. However, the phages that infect *A. baumannii* have a notoriously narrow host range, typically infecting only a small handful of strains, thus limiting their utility in phage therapy[7–9]. Furthermore, even when patients are treated with phage cocktails to which the bacteria is susceptible, resistance often develops rapidly[6,10].

The most important factor that drives host range is the compatibility between the phage receptor binding protein (RBP) and the bacterial receptor, typically a protein or molecule exposed on the cell surface such as lipopolysaccharide, type IV pili, or capsule[11,12]. In some cases, the bacterial capsule can serve a protective role, blocking access to receptors and providing phage resistance in organisms such as *Escherichia coli* and *Staphylococcus aureus*[13–15]. However, some bacteriophages are able to use capsule as a receptor, especially in species that are typically encapsulated such as *Klebsiella pneumoniae*[16]. Most *A. baumannii* phages characterized to date use the bacterial capsule as their receptor, with surprisingly few other receptors reported[4]. The *A. baumannii* capsule is a crucial virulence factor that protects the bacteria from various threats including killing by the eukaryotic host immune system as well as from small molecules like antibiotics[17–23]. The bacterial capsule is composed of capsular exopolysaccharides that come in a staggering number of variations in sugars and specific linkages, with over two hundred unique capsule loci producing over seventy distinct capsule structures in this organism[24]. Individual phages often carry a capsule-specific depolymerase associated with their tail spike protein that allows them to bypass this physical barrier[25]. Because of this narrow specificity, identifying phages for any given *A. baumannii* strain is challenging and bacteria quickly become resistant to capsule-dependent phages by modifying or losing their capsule both *in vitro* and *in vivo*[10,20,26,27]. A deeper understanding of the receptor usage of phages commonly used in phage therapy is crucial to circumventing bacterial resistance and rationally designing phage cocktails[28–30]. Despite the relatively low diversity of reported *Acinetobacter* phages, surprisingly little is understood about how these phages adsorb to their host. In this work we report the isolation of three lytic phages, StAb1, StAb2, and StAb3, that represent the three major morphological types of double-stranded DNA phages (podo-, myo-, and siphoviruses). StAb1 and StAb2 represent the two of the most commonly isolated groups of *Acinetobacter* phages, Friunaviruses and Twarogviruses based on genomes available on NCBI, and are closely related to phages previously used in phage therapy[10,31]. We find that StAb1 and StAb2 are both highly specific for a particular capsule type, but intriguingly, StAb2 is also able to infect any *A. baumannii* strain tested lacking a capsule. In contrast, StAb3 is highly unusual in its ability to infect most *A. baumannii* strains irrespective of capsule type. Here, by leveraging escape mutant analyses, we identify the receptors for each of these phages, revealing different strategies employed by *Acinetobacter* phages to overcome the capsule barrier. We find that rationally combining phages based on their distinct capsule interaction strategies can reduce development of bacterial resistance, demonstrating potentially therapeutically useful insights into phage cocktail selection.

## Results

### Isolation and classification of three *A. baumannii* phages

We isolated three *A. baumannii* phages from wastewater using two different *A. baumannii* clinical isolates hosts: StAb1 and StAb2 were isolated on 398 and StAb3 was isolated on MC47.2. All phages form clear plaques and exhibit lytic activity against their isolation host strain **(Fig 1A)**. Transmission electron microscopy (TEM) reveals that these are all tailed phages: StAb1 is a podophage with a head measuring ∼63 (±4.9) nm, StAb2 is a myophage with a hexagonal head of 78 (±4.6) nm width and 103 (±5.9) nm height, and StAb3 is a siphophage with a head about 75 (±2.4) nm in diameter, but with a notably long, non-contractile tail (336 ±8.4 nm) **(Fig 1B)**. We next sequenced the phage genomes and found that StAb1 has a genome of approximately 42 kb and is in the *Autographiviridae* family, genus *Friunavirus* (**Fig 1C and S1A Fig**). StAb2 has a genome of approximately 165 kb and is a member of the *Straboviridae* family, which also contains the model *E. coli* T4 phage; within this family it belongs to the *Acinetobacter* specific *Twarogvirinae* subfamily (**Fig 1C and S1B Fig**). Within this subfamily, StAb2 is a strain of *Lazarusvirus fhyacithree*, exhibiting >90% identity and substantial synteny to other phages of the *Lazarusvirus* genus, including a group of phages used in the Patterson phage therapy case (AC4, Navy1, Navy4, and Navy97) as well as two recently reported phages, DLP1 and DLP2, that were isolated on acapsular strains of *A. baumannii*[31]. StAb3 has a genome of approximately 79 kb and is a member of an unclassified taxon with similarity to three other taxonomically unclassified *Acinetobacter* phages described as having a broad host range, with 97% identity to EAb13, 67% identity to Mystique, and 68% identity to vB_AbaS_TCUP2199[32–34] (**Fig 1C and S1C-D Figs**). In spite of the relatively low nucleotide identity between some of these phages, they do exhibit substantial synteny and high identity at the protein level and are likely members of a shared, mostly unexplored taxon of *Acinetobacter* phages **(Fig 1C and S1D Fig**).

**Fig 1.**
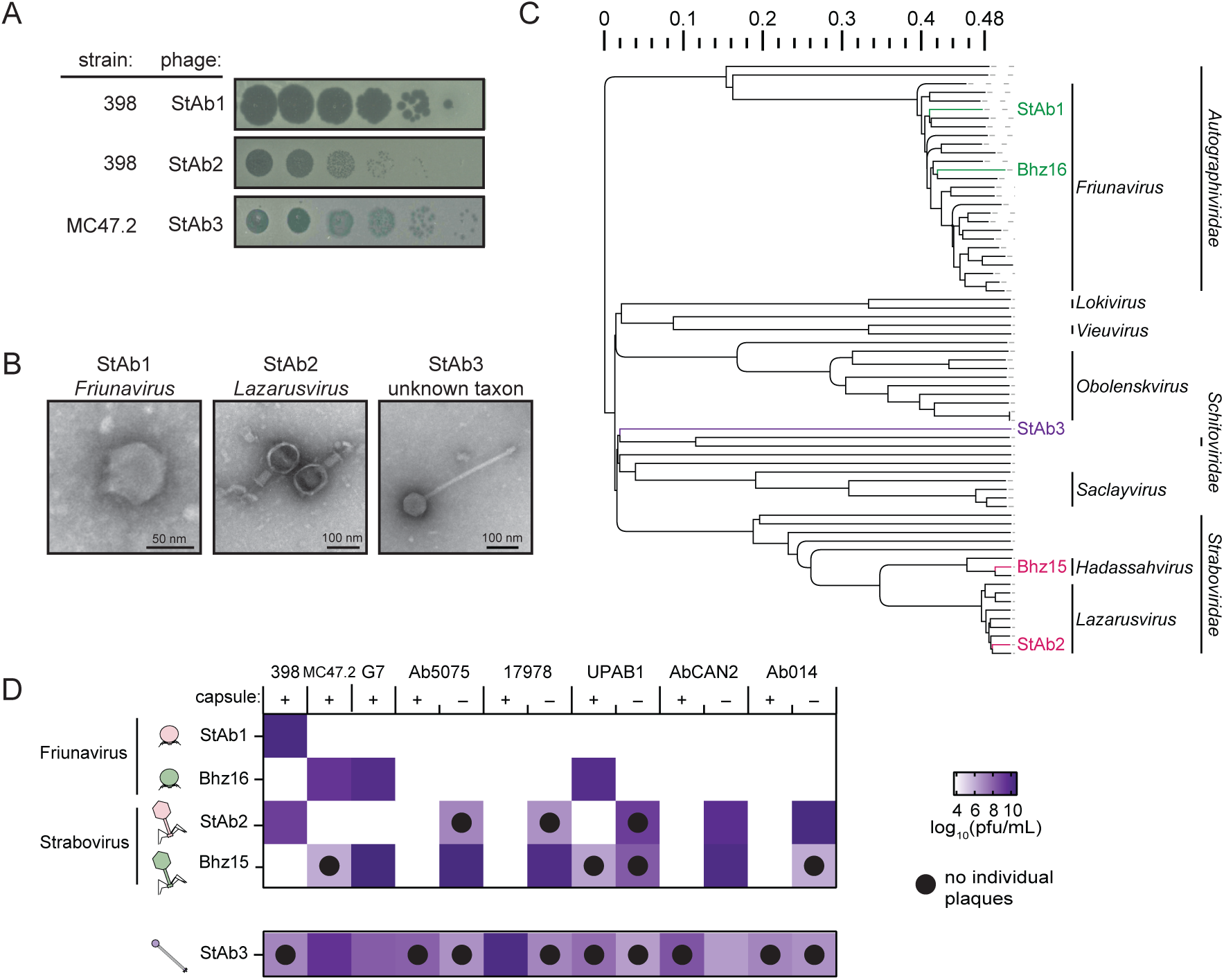
Isolation of three representative *A. baumannii* phages. A) Plaque assays of each phage on the indicated isolation host. B) TEM images of StAb1, StAb2, and StAb3. C) Phylogenetic tree of the 65 *Acinetobacter*-infecting phages with complete genomes available in RefSeq, with phages sequenced in this paper labeled. D) Host range analysis of StAb1, Bhz16, StAb2, Bhz15, and StAb3 on a panel of *A. baumannii* strains with and without capsule with colors indicating the average log_10_(PFU/mL) of three independent replicates. Phages were titered using double overlay plaque assays under optimal conditions for each phage, as described in Materials and Methods. Black dots indicate combinations where phages did not form individual plaques in at least 2/3 replicates.

We next determined the host range of these three phages across a panel of contemporary clinical isolates of *A. baumannii.* We observed that both StAb1 and StAb2 form plaques only on their isolation host, with no visible clearance or lysis visible on any other wild-type strains **(Fig 1D)**. This narrow host range is similar to most known *Acinetobacter* phages, which typically use the highly variable capsule as the receptor, and we hypothesized that the host range of StAb1 and StAb2 is likewise dependent on capsule type. In contrast, StAb3, while not able to form visible individual plaques on every strain, is able to form zones of clearance to high titers for each of the tested clinical isolates **(Fig 1D)**.

In the course of our host range experiments, we noted StAb2 could occasionally form plaques on one of the clinical isolates in our collection, Ab5075. Upon further investigation, we discovered that one of our glycerol stocks contained an Ab5075 variant that had spontaneously lost capsule, and that this capsule deficient strain is susceptible to StAb2 infection, while its wild-type counterpart is fully resistant. This surprising observation suggests that, in the absence of capsule, StAb2 may have a broader host range and that capsule is not a requirement for infection of StAb2, unlike many other reported *Acinetobacter* phages. We previously showed that a mutant in the initiating glycosyltransferase, PglC, in *A. baumannii* strain 17978 was unable to produce capsule and that this could be complemented by supplying *pglC* in a plasmid[35]. We confirmed these phenotypes by employing a previously validated density gradient assay[36–38] **(S2A-B Figs).** Bacteria with more capsule have a lower density, while bacteria with decreased or absent capsule have a higher density and thus travel further down the density gradient upon centrifugation[36–38]. Consistent with StAb2 infecting acapsular strains more broadly, we found that the *pglC* mutant is sensitive to StAb2, while resistance was restored in the complemented strain **(S2C Figs)**. We assembled a panel of clinical isolates which had lost their capsule either through targeted deletions or spontaneous mutations and validated the loss of capsule with both sequencing and the density gradient (**S2 Fig and S1 Table**). We found that, consistent with most other reported *A. baumannii*-infecting phages, StAb1 retained its narrow host range, only forming plaques on its isolation strain, 398 **(Fig 1D)**. However, StAb2 was able to infect capsule-deficient variants of five distinct clinical isolates of *A. baumannii* **(Fig 1D),** demonstrating that StAb2 has a surprisingly broad host range in the absence of capsule, implying that capsule functions as a barrier to infection by this phage. Finally, both the wild-type strains and capsule mutants are susceptible to StAb3, though in most cases the capsule mutants do show reduced sensitivity compared to the wild-type strains **(Fig 1D)**. Together, these results demonstrate that phylogenetically diverse *Acinetobacter* phages exhibit key differences in their interactions with capsule, which can impede or promote replication of distinct phages.

### *Acinetobacter* phage StAb1 requires capsule for infection

To systematically determine the key bacterial factors that contribute to phage host range, we first set out to analyze bacterial clones that had acquired resistance to StAb1. For StAb1 planktonic infection, we typically observed an initial period during which no bacterial growth is detected, followed by an increase in optical density, suggesting the presence of escape mutants (**Fig 2A**). Clones of the surviving bacteria were isolated and durable phage resistance was confirmed (i.e. no plaques observed in double overlay assays).

**Fig 2.**
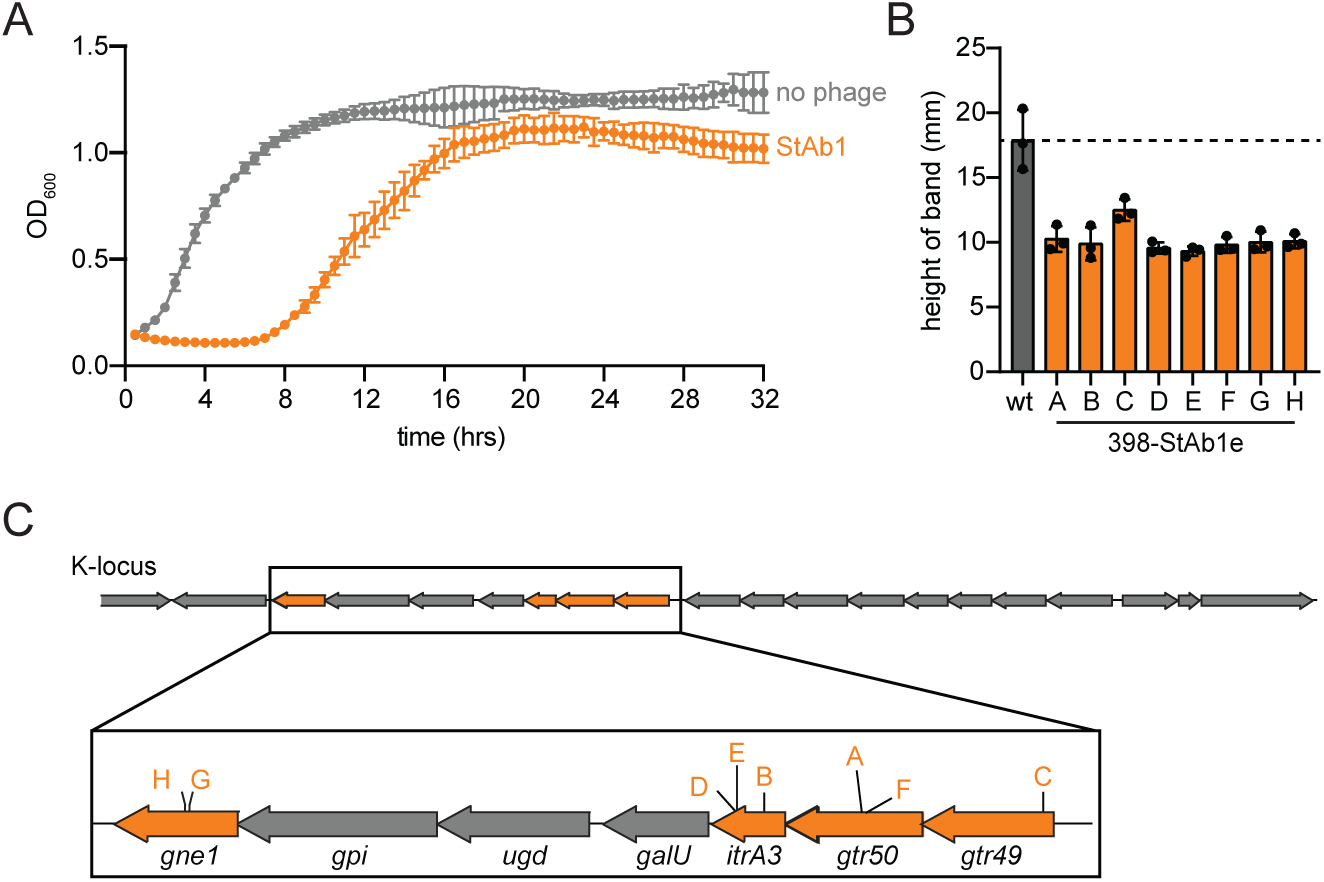
*A. baumannii* escapes StAb1 by losing its capsule. A) Bacterial growth monitored via absorbance at OD_600_ with and without StAb1 phage. The average of three technical replicates is presented, error bars representing standard deviation (sd). B) StAb1 phage escape mutant capsule formation assessed via silica-based density gradient experiments and compared with the wild-type (wt) parental strain 398 (gray). Three independent replicates and their average is presented with error bars representing sd. C) 398 capsule biosynthesis operon (K-locus) indicating the location of SNPs for each 398-StAb1 escape mutant corresponding to strains in (B).

We assessed capsule production in these escape mutants using the density gradient assay and determined that all StAb1 escape mutants likely lost or produce dramatically less capsule **(Fig 2B)**. Consistent with these results, 398-StAb1 escape mutants all have mutations in genes found in the K-locus, the locus encoding most genes required for capsule synthesis **(Fig 2C and S2 Table)**[24]. To extend these findings to the larger *Friunavirus* genus, we acquired a closely related phage, Bhz16, and found that while it has specificity for a different capsule type, the pattern is largely the same. Bhz16 can form plaques only on three strains predicted to have a similar capsule, UPAB1, MC47.2, and G7, but is unable to replicate on an acapsular derivative of UPAB1 or any other acapsular strain (**Fig 1C and S3 Table**)[24,39]. As with StAb1, G7-Bhz16 escape mutants have reductions in capsule and mutations in the K-locus (**S2E Fig and S2 Table)**. This requirement for capsule likely drives the very limited host range of *Friunavirus* phages, with only capsule types complementary to each phage able to support infection.

### Bacteria can become resistant to StAb2 by increasing or altering capsule

We also obtained isolates of the *A. baumannii* 398 strain that were resistant to StAb2 (**Fig 3A**). Sequencing revealed a more diverse suite of mutations in StAb2 resistant strains than those found for StAb1. One escape mutant, 398-StAb2eA, mapped to the K-locus **(Fig 3B and S2 Table)**. We noted that when grown on an agar plate, this strain is more mucoid than the wild type, and its migration in the density assays suggests increased capsule production (**Fig 3C**). This strain has a point mutation in *wzc*, which encodes a tyrosine kinase that plays a key role in regulation of capsular polysaccharide synthesis and transport across the membrane[40]. Specific point mutations in *wzc* have been previously shown to result in overproduction of capsular polysaccharides[23,41]. All other StAb2 escape mutants have single nucleotide polymorphisms (SNPs) in genes associated with lipooligosaccharide (LOS) synthesis. These include mutations in A1S_2903, a gene involved in an early step of LOS synthesis[42]. We therefore hypothesized that LOS was essential for StAb2 infection, potentially serving as a receptor. However, contrary to this hypothesis, we found no change in susceptibility for a Δ*lpsB* mutant to StAb2 **(Fig 3D)**. It was previously shown that ΔA1S_2903 and Δ*lpsB* synthesize the same truncated LOS, indicating that changes in LOS do not drive StAb2 resistance in these strains and that LOS is not the receptor for StAb2[42]. Density gradient assays of the StAb2 escape mutants suggest that these strains have equal or increased capsule production (**Fig 3C**). This, together with the reduced sensitivity of these strains to StAb2 suggest potential unexplored crosstalk between LOS and capsule synthesis in *A. baumannii* that should be investigated further in the future. In conclusion, in contrast to StAb1, StAb2 escape mutants are not deficient in capsule synthesis, and instead likely acquire resistance through increased or altered capsule. Importantly, some form of capsule, possibly altered, is maintained, consistent with our observations that acapsular strains are susceptible to StAb2 infection. Together, these data demonstrate that the capsule provides a critical first line of defense against StAb2.

**Fig 3.**
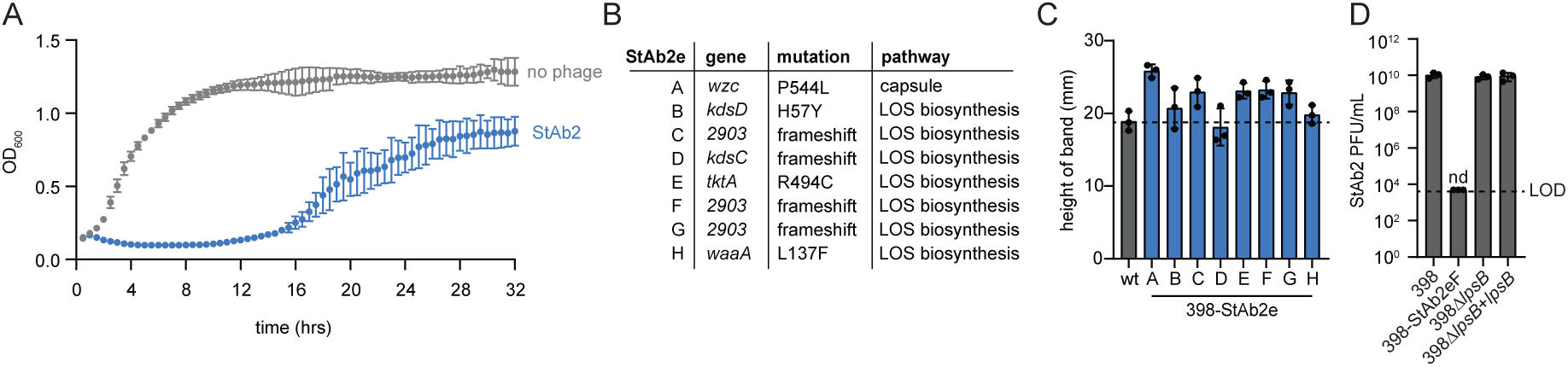
*A. baumannii* evades StAb2 by altering or increasing capsule. A) Bacterial growth monitored via absorbance at OD_600_ rebounds after StAb2 phage addition, suggesting bacterial escape mutants arise rapidly. The average of three technical replicates is presented with error bars representing sd. B) Summary of mutations identified in 398-StAb2 escape mutants. C) StAb2 phage escape mutant capsule formation assessed via silica-based density gradient experiments and compared with the wild-type (wt) parental strain 398 (gray). D) Quantification of plaque assays with StAb2 on the indicated strains. Limit of detection (LOD) = 5 x 10^3^ PFU/mL. nd: not detected.

### *Twarogvirinae* phages use an outer membrane porin, CarO, as an alternate receptor

To identify the non-capsular receptor for StAb2 we employed a transposon mutagenesis-based screening approach. We generated a high-density transposon mutant library of one of the StAb2-sensitive, capsule-deficient strains, UPAB1 Δ*wzy,* challenged the library with StAb2, and isolated surviving mutants. Phage resistance was verified by plaque assays, and we then identified the transposon insertion sites by sequencing. Three resistant mutants contained transposon insertions in the gene encoding the outer membrane porin, CarO (Carbapenem-associated outer membrane protein), suggesting that CarO may be the receptor for this phage **(Fig 4A)**. Additionally, we generated escape mutants to StAb2 in the acapsular 17978 Δ*pglC* strain and found that each clone has a SNP in the *carO* gene. Four of these candidates have a SNP at position 110 (escape type T) in the *carO* nucleotide sequence, resulting in a premature stop codon and truncation of the CarO protein, while the remaining four have a SNP at position 638 (-G) (escape type F), resulting in a frameshift mutation **(Fig 4A and S2 Table)**. CarO is a conserved porin involved in nutrient uptake and possibly facilitates carbapenem entry into the bacterial cell [43–45]. While the gene *carO* is interrupted or absent in some *A. baumannii* isolates, it is present in most *A. baumannii* genomes, explaining why StAb2 can infect any of the capsule-deficient strains tested[44,46,47].

**Fig 4.**
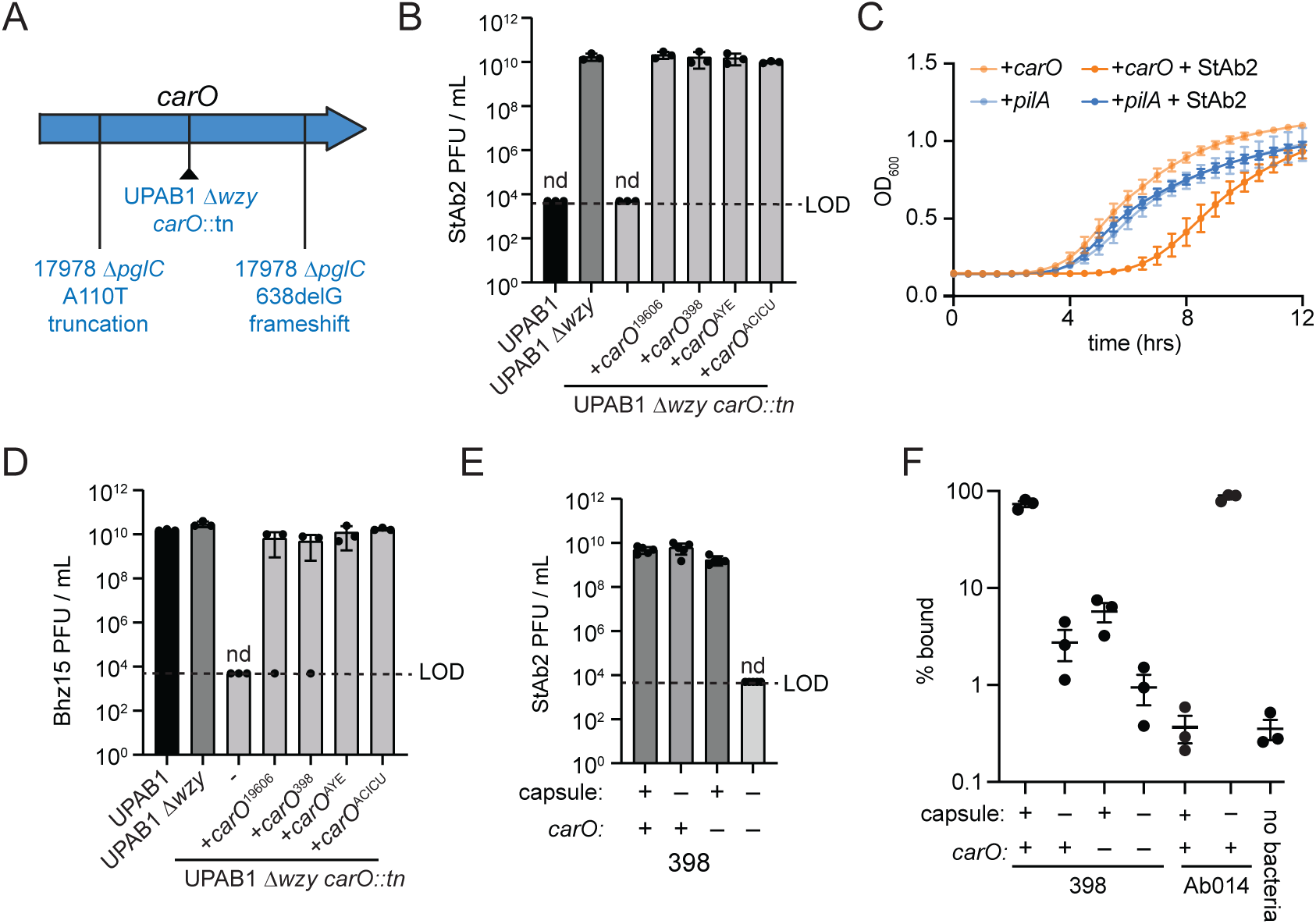
CarO is an alternate receptor for StAb2 in the absence of capsule. A) Mutations identified in CarO for StAb2 resistant strains: 17978 Δ*pglC*-StAb2eA-H and UPAB1Δ*wzy carO*:tn, as indicated in the schematic. B) Quantification of plaque assays with StAb2 on the indicated UPAB1 strains and mutants complemented with *carO* from the indicated strain. C) Growth of *E. coli* expressing either CarO or PilA (control) with or without addition of StAb2 at MOI = 1000. The average of three independent replicates each with three technical replicates is presented with error bars representing sd. D) Quantification of plaque assays with Bhz15 on the indicated UPAB1 strains ectopically expressing different *carO* alleles, as in (B). E) Quantification of plaque assays with StAb2 on the indicated strains: 398 (capsule+, CarO+), 398-StAb1eA (capsule-, CarO+), 398 Δ*carO* (capsule+, CarO-), or 398-StAb1eA Δ*carO* (capsule-, CarO-). F) Adsorption of StAb2 after 10 min of incubation with the 398 strains from (E), as well as Ab014 (capsule +) and Ab014-cm (capsule –). B, D-F) LOD = 5 x 10^3^. nd = not detected. Three independent replicates and their average is presented with error bars representing sd.

We identified and cloned four CarO proteins from different *A. baumannii* strains representing different identified subtypes, and found that all were able to complement the transposon insertion strain and restore phage replication when expressed ectopically **(Fig 4B)**[47]. Further validating that CarO serves as a receptor for StAb2, we also found that when CarO^19606^ is expressed in *Escherichia coli*, StAb2 was able to inhibit growth in planktonic infections. We did not observe this StAb2-dependent growth defect in *E. coli* expressing PilA, which was employed as a control **(Fig 4C)**. Together, these experiments demonstrate that for an acapsular *A. baumannii*, CarO is necessary for StAb2 infection, and that expression of CarO is sufficient for phage infection in even a distantly related bacterial species.

We hypothesized that *Twarogvirinae* carry a conserved CarO-binding protein in addition to a unique capsule-binding (and/or degrading) protein specific for the capsule type of its host. Supporting this idea is the fact that *carO* was identified in an escape mutant analysis of a related *Lazarusvirus*, Navy-1, following *in vitro* selection, though the authors did not consider a role for *carO* as a potential phage receptor[10]. These authors also noted mutations in the K-locus in resistant bacteria arising both *in vivo* and *in vitro,* though whether these strains had lost capsule was never tested. Further, CarO was also identified as the receptor for the *Lazarusvirus* DLP2, a phage isolated on an acapsular strain[31]. To test whether the capsule/CarO dual receptor usage is similar for another *Twarogvirinae* phage of a different genus, we obtained Bhz15, which genome sequencing revealed is, like StAb2, a *Strabovirus* of subfamily *Twarogvirinae*, but of the genus *Hadassahvirus*. Bhz15 has a narrow host range, infecting only its host, G7, and like Bhz16, which was isolated on the same host strain, Bhz15 is also able to infect the UPAB1 strain with a similar or identical capsule type **(S3 Table)**. Much like StAb2, Bhz15 can infect all acapsular *A. baumannii* strains **(Fig 1C)**. Bhz15 readily infects the capsule mutant UPAB1 Δ*wzy*, but was not able to infect UPAB1 Δ*wzy carO*::tn, a mutant resistant to StAb2 with a transposon interrupting CarO **(Fig 4D).** Further, the ability of Bhz15 to infect the CarO transposon mutant was restored upon complementation of CarO **(Fig 4D)**. This demonstrates that Bhz15, like StAb2, is prevented from accessing CarO by most capsule types, and in the absence of capsule, can use CarO as a receptor, suggesting that this receptor usage pattern is a feature of the *Twarogvirinae*.

### StAb2 can use either CarO or cognate capsule for cell entry

While CarO is clearly essential for infection in non-isolation hosts and sufficient for infection of entirely distinct bacteria, StAb2 and Bhz15 can each replicate on their capsule-proficient isolation strains, 398 and G7, respectively. In these strains, CarO is likely cloaked by the presence of capsule, implying these phages utilize the capsule or another external structure as a primary receptor. We wondered if CarO was required for infection of 398 by StAb2, or if the capsule could serve as the sole receptor. To test this, we generated a *carO* deletion in both the wild-type 398 and one of the capsule-deficient 398 StAb1 escape mutants (398-StAb1eA) and tested StAb2 infectivity and adsorption with these strains. StAb2 has a similar efficiency of plating (EOP) on 398-StAb1eA, which lacks capsule, compared with Δ*carO* and the wild-type 398. **(Fig 4E)**. The strain lacking both capsule and CarO, 398-StAb1eA Δ*carO*, is resistant to StAb2, demonstrating that StAb2 requires either CarO or capsule for infection **(Fig 4E)**. Indeed, direct measurement of adsorption revealed that about 80% of the bacteriophages adsorb to wild-type 398 within 10 minutes, about 10% can adsorb to 398 bearing only intact capsule or CarO, while phage adsorption to a strain lacking both receptors is below the limit of detection **(Fig 4F)**. Together, this demonstrates that StAb2 can use either CarO or capsule in infecting 398, though adsorption is considerably improved by having both.

To assess if the inability of StAb2 to infect most other capsulated *A. baumannii* strains is due to an inability to adsorb or to penetrate the cell wall, we assessed the ability of StAb2 to adsorb to wild-type clinical isolate Ab014 and its capsule mutant counterpart. We found that StAb2 is entirely unable to adsorb to the wild type, but after 10 minutes, close to 100% of phage particles have adsorbed to the Ab014 capsule mutant **(Fig 4G)**. This demonstrates that while StAb2 can adsorb to its cognate capsule type found in 398, it is unable to adsorb to a strain with a non-cognate capsule type, Ab014. Further, this demonstrates that the *A. baumannii* capsule does indeed prevent StAb2 from accessing its receptor, CarO. StAb2 adsorption to CarO^Ab014^ is also notably stronger than to CarO^398^, though the weaker adsorption to CarO^398^ does not meaningfully impact phage replication **(Fig 4E)**.

While Bhz15 and StAb2 can both infect acapsular *A. baumannii* strains using CarO as a receptor, their host range amongst capsular *A. baumannii* strains is distinct. The ability of phages to penetrate extracellular polysaccharides such as capsule or biofilms is often dependent on the presence of phage-encoded depolymerases[25,48,49]. Analysis of the proteins encoded by StAb2 using two machine learning tools that predict phage depolymerases identified CDS_0239, annotated as a tail fiber protein, as the most likely depolymerase (**S3A Fig**)[50,51]. Characterized depolymerases in *Acinetobacter*-infecting Friunaviruses are similarly tail spike proteins and are encoded directly before lysins, as is CDS_0239[25]. A closer examination of the tail fiber genes revealed that, while most genes are very similar within the *Lazarusvirus* and *Hadassahvirus* genera, homologs of CDS_0239 exhibit substantial variation that is inconsistent with broader phylogeny, reminiscent of the hypervariable regions found within the *A. baumannii* K-locus **(S3A-C Fig)**[24,52]. A multiple sequence alignment of this gene with its homologs from other annotated Lazarusviruses and Hadassahviruses revealed considerable mosaicism, with the last ∼250 amino acids exhibiting the highest variability of this gene (**S3D Fig**). This C-terminal domain varies substantially between closely related phages and does not track with phylogeny, suggesting it is under strong positive selection, similar to *Friunavirus* tail spike proteins, wherein the N-terminal portion is highly conserved and the divergent C-terminal domains have been proposed to determine specificity for capsular polysaccharides (**S3D Fig**)[25]. The second variable region of this protein (amino acids 69-976) contains the previously identified pyocin knob domains which are variable between *Lazarusvirus* and *Hadassahvirus* strains and are proposed to be involved in receptor interactions[53].

Based on these analyses and our experimental observations, *Twarogvirinae* phages StAb2 and Bhz15 attach to and likely degrade the capsule of susceptible hosts using the highly variable long tail fiber protein, and then use a second, conserved receptor-binding protein to bind CarO. Thus, the *A. baumannii* capsule blocks access to CarO from the vast majority of Twarogviruses except for the small subset with a depolymerase capable of degrading its particular capsule type (e.g. StAb2 for 398 capsule; Bhz15 for G7 capsule). However, they can infect acapsular *A. baumannii* strains as the CarO protein is exposed.

### StAb3 requires an uncharacterized cell surface polysaccharide for infection

In contrast to StAb1 and StAb2, StAb3 is able to infect a wide range of *A. baumannii* strains, including both capsular and acapsular strains. It seemed unlikely that a depolymerase could target multiple, diverse capsule types, as they are often composed of entirely unique polysaccharides, and further, StAb3 could infect entirely acapsular strains as well. To investigate how StAb3 is able to infect diverse capsule types, we obtained MC47.2 escape mutants from StAb3. All of these escape mutants were comparable to the wild type in the density gradient assay, suggesting that the capsule is unchanged **(Fig 5A)**. All isolates also have two IS4 family transposase genes, also found in another section of the wild-type chromosome, inserted in the gene MLMLPJ_10955 **(Fig 5B and S2 Table)**. This gene is homologous to BcsB, a protein that in other bacterial species is involved in the synthesis of cellulose[54–57] (**Fig 5C, S4 Fig, and S4 Table**). BcsB is encoded in a locus that also contains homologs to genes involved in the synthesis of other exopolysaccharides, including Poly-N-acetylglucosamine (PNAG)[57]. This locus includes a putative outer membrane porin to facilitate polysaccharide transport across the outer membrane, a putative diguanylate cyclase which synthesizes cyclic-di-GMP, an important regulator of multiple exopolysaccharide transport systems, and a structural homolog to the C-terminus of PgaB, involved in PNAG binding during its synthesis[57,58]. The locus also includes a homolog of WecB, or UDP-N-acetylglucosamine-2-epimerase, which synthesizes UDP-N-acetylmannosamine (ManNAc). Further, a StAb3 escape mutant we isolated from a different parental *A. baumannii* strain, Up280, carries a single SNP within this *wecB* gene **(Fig 5A and S2 Table)**[58,59]. Previous studies identified a similar operon in *E. coli* as producing a sugar essential for the bacteriophage N4 to adsorb to and infect *E. coli*[60,61]. The authors proposed that these genes encode an exopolysaccharide transport system producing a ManNAc-based polysaccharide that serves as the receptor for N4[60,61]. Similarly, we propose that this locus produces a still uncharacterized ManNAc-containing exopolysaccharide that serves as a receptor for StAb3 **(Fig 5C)**. This polysaccharide likely protrudes beyond the capsule, providing StAb3 with a point of attachment even when capsule is present and enabling it to attach to and infect *A. baumannii* independent of capsule. We note that infection is in some cases diminished when capsule is removed (**Fig 1C**); one potential explanation for this phenotype is that the capsule may help orient the phage and/or sugar, and when it is absent infection proceeds with a reduced efficiency.

**Fig 5.**
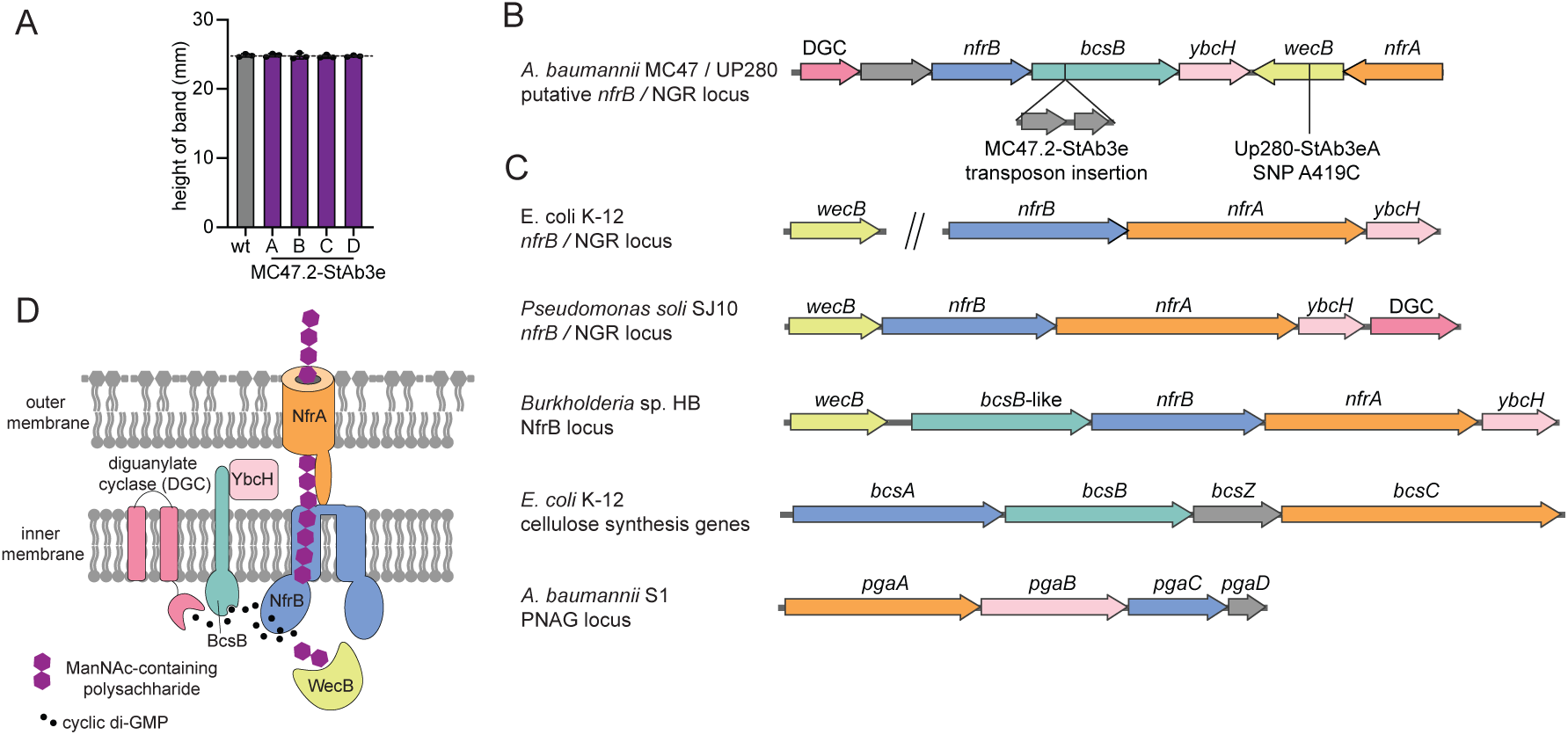
Escape mutants to StAb3 arise in a predicated EPS biosynthesis and secretion pathway. A) StAb3 phage escape mutant capsule formation assessed via silica-based density gradient experiments and compared with the wild-type (wt) parental strain MC47.2 (gray). Three independent replicates and their average is presented with error bars representing sd. B) Locus containing a transposon insertion in MC47.2 and Up280 escape mutants to StAb3 infection. Shared predicted function and homology to related EPS export pathways is indicated by color, see S4 Fig and S4 Table. C) Model of predicted exopolysaccharide secretion system based upon the structural homology and localization of proteins in related EPS secretion systems.

### Phages can be paired to counteract *A. baumannii* resistance

We have identified capsule-dependent (StAb1 and Bhz15) and capsule-restricted (StAb2 and Bhz16) *A. baumannii* phages. These findings suggest an intriguing possibility: capsule-restricted phages should be able to suppress the growth of capsule-deficient escape mutants that emerge rapidly when bacteria encounter capsule-dependent phages, such as StAb1, and extend the efficacy of these phages in combination. Supporting this concept, escape mutants to StAb1 remain susceptible to StAb2, and most escape mutants from StAb2 remain susceptible to StAb1 **(Fig 6A-B)**. Further, we found that when 398 (for StAb1 and StAb2) or G7 (for Bhz15 or Bhz16) is treated with a single phage, escape mutants arise within 20 hours and expand rapidly (**Fig 6C-D and S5A-B Fig**). However, the combination of StAb1 and StAb2 on 398 or Bhz15 and Bhz16 on G7 fully suppress emergence of escape mutants throughout the entire course of our experiments, 36 hours (**Fig 6C-D** and **S5A-B**).

**Fig 6.**
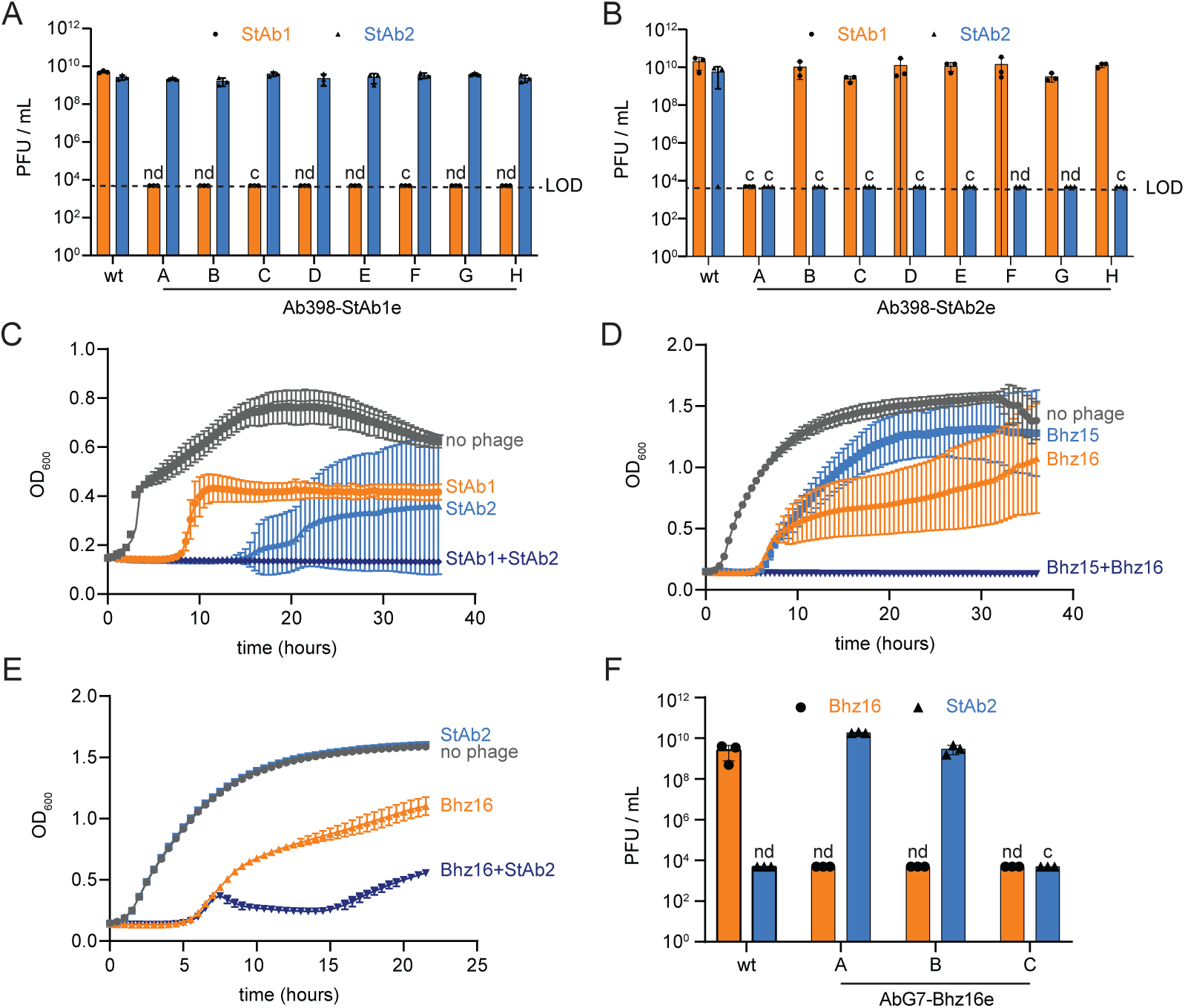
Combinations of phages prevent or delay bacterial resistance. A-B) Quantification of plaque assay with StAb1 and StAb2 on wild-type 398 and 398 StAb1 (A) and 398 StAb2 escape mutants (B). LOD = 5 x 10^3^. nd = not detected. c = areas of clearing, but no individual plaques, observed. C-E) Growth of 398 (C) or G7 (D-E) with the indicated individual phages or phages combined. Phages were added to an MOI of 1, either individually or combined. The average of six (C&D) or three (E) technical replicates is presented with error bars representing sd. See figure S5 for two additional biological replicates. F) Replication of Bhz16 and StAb2 on either wild-type G7 or G7 Bhz16 escape mutants as in (A) and (B).

One major hurdle with phage therapy is the need to either test many phages to identify one that can kill the bacteria causing the infection or even to isolate phages on that exact strain, due to their typically exquisite host-range specificity. However, our findings that Twarogviruses StAb2 and Bhz15 can infect a broad range of acapsular strains suggests that these phages could be used as a secondary phage in a cocktail against many *Acinetobacter* strains as long as a single capsule-dependent phage can be found that infects that strain, reducing labor required to isolate multiple strain-specific phages. We thus reasoned that StAb2 or Bhz15 should suppress escape mutants from arising in a non-native host strain when paired with a strain-specific, capsule-dependent phage that drives the emergence of acapsular escape mutants. To test this, we first combined Bhz15 with StAb1 to kill 398, which Bhz15 cannot typically infect, and found that the addition of Bhz15 was not able to prevent bacterial resistance from arising (**S5C Fig**). This is possibly due to a 398 host-specific factor such as a bacteriophage defense system that restricts Bhz15. However, we further combined StAb2 with Bhz16 on G7, which StAb2 cannot typically infect. While it did not fully prevent bacterial resistance from arising, it did delay and reduce escape mutants **(Fig 6E and S5D Fig)**. We also tested the resistant bacterial isolates following Bhz16 infection and found that two out of the three were susceptible to StAb2 **(Fig 6F)**, suggesting that these phages could also function effectively if given sequentially rather than in combination. In summary, these data demonstrate that bacterial resistance can be eliminated or reduced by pairing a capsule-dependent phage with a *Strabovirus* that uses CarO as its receptor.

## Discussion

Capsule constitutes a formidable barrier that protects bacteria from diverse environmental stressors, including phages. In this work we describe three mechanisms by which diverse phages interact with or bypass this physical barrier (**Fig 7**). The selective pressure imposed by phage infection likely plays a role in driving the extensive diversification of the capsule. *A. baumannii* produce more than seventy types of capsule, which provides the first immunity barrier against the phages[24]. Phages like StAb1 and StAb2 overcome this defense by attaching to and degrading the capsule to access the cell surface. While this seems to be necessary and sufficient for StAb1 infection, StAb2 and other *Twarogvirinae* phages can also use CarO as a receptor, which is typically occluded by capsule. Both capsule-specific phages encode predicted depolymerases that are highly variable even within very close relatives[62]. The diversity of *A. baumannii* capsular structures is likely matched by specific and diverse depolymerases in these phages, which results in an effective phage attack with a narrow host range. A less well-characterized siphophage, StAb3, appears to employ an alternative strategy. Our experiments strongly suggest that this phage, which has a notably long tail, instead recognizes a yet to be characterized ManNAc-containing glycan, somehow being able to inject its DNA without depolymerizing the capsule.

**Fig 7.**
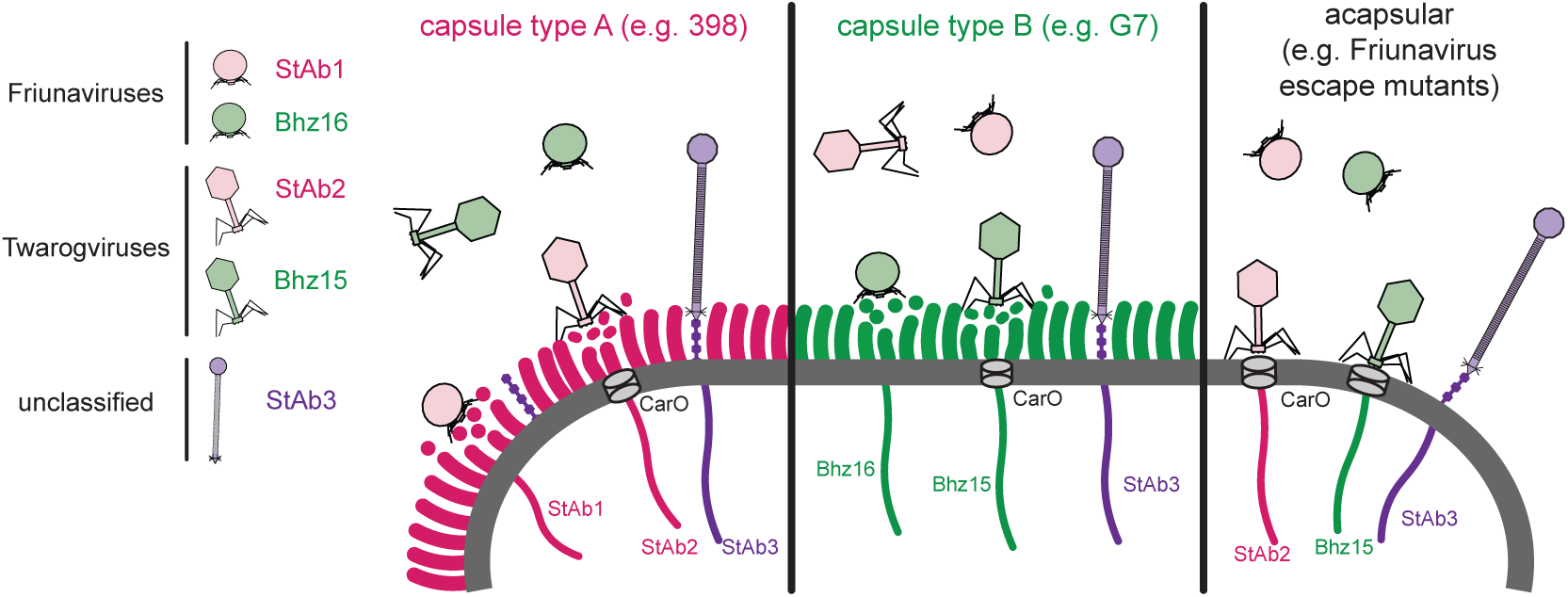
Model of bacteriophage-capsule interactions. Model for *Friunavirus* (StAb1-like), *Twarogvirinae* (StAb2-like), and StAb3-like phage interaction with *A. baumannii* capsule.

Our study demonstrates the importance of phenotypically characterizing escape mutants, in addition to identifying genetic changes. Prior studies generally assumed that mutations in the K-locus are indicative of capsule loss; however, we find that one mutation located in the K-locus that led to StAb2 resistance has an increase in capsule production, rather than capsule loss, shown by our density-gradient assay. Furthermore, when we isolated 398 StAb2 escape mutants we found that most had SNPs in genes related to LOS synthesis, including A1S_2903. At first glance, this would suggest that StAb2 might use LOS as a receptor. However, a *lpsB* mutant, which in other strains has the same truncated LOS as a A1S_2903 mutant, remains susceptible to the phage, ruling out a role for LOS as the receptor[42]. Previous research demonstrated that while A1S_2903 and lpsB mutants have the same LOS structure, they are phenotypically distinct, with A1S_2903 mutants having a more pronounced growth defect, increased sensitivity to cell-wall targeting antibiotics, vancomycin and bacitracin, and increased propensity to form chains and clusters[42]. Many of the escape mutants we isolated, including those with SNPs in A1S_2903, migrate differently in the density gradient assay, suggesting increased or altered capsule. Thus, phenotypic characterization demonstrates mutations in the LOS biosynthesis pathway may affect capsule, revealing unexpected interactions between the LOS and capsule biosynthesis pathways. Our work shows that phages exert unique selective pressures that can lead to changes in bacterial physiology, revealed by escape mutant analyses, that expose knowledge gaps about basic bacterial processes, such as the unexpected cross-talks between LOS and capsule synthesis. More research is needed to understand how these pathways are interconnected and impact the assembly of the capsule.

CarO has been previously associated with resistance to *Twarogvirinae* phages. In a genomic analysis of the phages used for therapy in the Patterson case, an escape mutant with resistance to phage AB-Navy1 was isolated and has a mutation in *carO*; however, the authors proposed that CarO might be changing the cell wall and did not discuss the possibility that CarO might be acting as receptor[10]. It is not clear why this mutation in CarO alone would be sufficient to abolish phage infection unless the capsule in this strain is not able to be used as a receptor and is not masking CarO to the same extent. Perhaps the strain produces less capsule *in vitro* due to the lower selective pressure for capsule, while *in vivo*, capsule production is higher and thus losing CarO alone would never be sufficient to lead to full resistance. Without a phenotypic analysis of capsule, it is impossible to distinguish between these possibilities. It was notable that CarO mutations were not identified *in vivo* in the Patterson case, consistent with reports that CarO may play a role in inhibiting host immune response and facilitating adherence to and invasion into cells[44,63–65]. In another report, CarO was identified as the receptor of the DLP2 phage, which is related to StAb2[31]. In that work, the authors identified CarO through a transposon screening in a capsule deficient mutant[31]. However, the role of CarO in capsulated strains was not investigated. Since multiple Twarogviruses can interact with CarO, we speculate that the CarO binding protein is present in all of these phages and is less variable than the capsule-specific binding protein.

StAb3, unlike many other *A. baumannii* phages, is able to infect strains from a broad range of capsule types. We found that StAb3 was unable to infect strains with an insertion or mutation in genes predicted to synthesize a ManNAc-containing exopolysaccharide in *A. baumannii*. It is tempting to speculate that StAb3 binds this exopolysaccharide, and facilitated by its long tail and predicted depolymerase, uses it as a guide through the capsule to reach the cell membrane. Interestingly, although the machinery for its synthesis is conserved among *A. baumannii* clinical isolates, the role and structure of this glycan remains unexplored. Further, a similar locus encoding an uncharacterized ManNAc-based polysaccharide serves as the receptor for N4 phage in *E. coli* [60,61]. More work is required to elucidate if these loci, which are also present in other species, are responsible for the synthesis of the same or different ManNAc-containing polysaccharides and to characterize its role in *A. baumannii* lifestyle and virulence. The StAb3 genome has over 65% similarity to that of unclassified phages EAb13, Mystique, and vB_AbaS_TCUP2199, all of which have been characterized as having a broad host range[32–34]. Further, both Mystique and StAb3 display impaired ability, but not complete inability, to infect acapsular strains. Several hypotheses can account for the impaired infection of StAb3 to acapsular strains. It is possible that the capsule stabilizes the phage tail in the proper conformation to enable injection of DNA. Alternatively, cross talks between capsule and exopolysaccharide receptors may occur. More work is needed to understand if our findings apply to other similar phages and how this exopolysaccharide interacts with bacterial capsule and phage infection.

Many phage therapy cocktails are assembled without knowledge of receptor usage of the phages included in the cocktail, leading to rapid resistance emerging in the bacterial host. There is increasing interest in rational design of phage cocktails using complementary phages with distinct receptors to target bacteria such as *Klebsiella pneumonia* and *Pseudomonas aeruginosa*[28–30]. We are furthering this effort with *A. baumannii* phages, identifying capsule-dependent and capsule-restricted phages, that when combined, can prevent or delay the emergence of escape mutants. This highlights the value in identifying phage receptors and applying this information to the rational design of phage cocktails. While several combinations of phages reduce bacterial resistance, not all combinations are equally or consistently effective, which demonstrate that there are still many nuances to phage infection that remain to be discovered. We speculate that cellular phage defense pathways also play a role in some cases. It is also likely that some of these phages may carry additional receptor binding proteins. Our detailed characterization of receptor usage will now open the door to rational design of phage therapy cocktails that include phages with different receptors, thereby preventing the rapid emergence of bacterial escape mutants.

## Materials and Methods

### Bacterial strains and growth conditions

Bacterial strains used in this study are listed in S5 Table. Unless otherwise noted, strains were grown on LB-agar plates containing LB broth (Fischer BioReagents, BP1427-2) and 1.5% agar (Fisher Bioreagents, BP1423-2) at 37°C. Colonies from these plates were used to inoculate cultures in LB broth grow at 37°C and shaking at 200rpm. When appropriate, LB broth and LB-agar plates were supplemented with 10 µg/mL chloramphenicol, 50 µg /mL apramycin, 100 µg/mL ampicillin, 50 µg/mL kanamycin, 50 µg/mL zeocin, and/or 300 µg/mL hygromycin B.

### Phage isolation

StAb1, StAb2, and StAb3 were isolated from wastewater based upon an adaptation of previously used methods[66]. In brief, wastewater samples were centrifuged at 2,300 x g for 8 minutes at 4°C, then supernatants were filtered through a Millex GS 0.22µm filter (Merck Millipore, SLGSR33SB) to remove any bacteria and particulate matter. Potential bacteriophages were enriched by combining 900µL of 2x LB broth, 100µL of an overnight culture of the isolation host bacterium, and 500µL of the filtered wastewater sample and incubating with shaking overnight at 37°C. Enrichments were centrifuged at 1,700 x g and filtered through a 0.22µm filter to remove any bacteria. 5-10µL of each sample was spotted onto a top agar plate prepared as described above for plaque assays. The bacteriophage titer for any samples which resulted in clearance of bacteria was determined utilizing the plaque assay described above. To ensure the isolate contained only one bacteriophage, a 10µL pipette tip was stabbed into an isolated plaque, swirled in a mixture of 180 µL of LB broth + 20 µL of an overnight culture of the isolation host bacterium and incubated with shaking at 37°C for approximately 6 hours. This sample was centrifuged and filtered to remove any bacteria as described below and plated on top agar plates with the isolation host bacterium. The isolation process was repeated for a total of three times to ensure isolation of a single clonal bacteriophage.

### Phage propagation

All phage strains utilized in this study are listed in S6 Table along with details about their isolation and propagation host bacteria. To propagate the phages, 10 mL of LB broth (with 10 mM MgCl_2_ for propagation of StAb2) was inoculated with 100 µL of an overnight culture of the propagation host bacterium and grown shaking until the OD_600_ reached approximately 0.4 at 37°C. Then, 100 µL of the previous phage stock was added to this culture (at an MOI of approximately 0.1) and incubation continued. Following clearing, samples were centrifuged at 6,200 x g for 8 minutes and the supernatant was filtered through a 0.22 µm filter to remove any remaining bacteria. Bacteriophage lysates were stored at 4°C for utilization in future experiments.

### Plaque assays

Double layer plaque assays were used to determine bacteriophage titers as well as assess the capacity of bacteriophages to infect specific hosts. A top agar was prepared with 4 mL of molten LB with 0.4% agar combined with 25-200 µL of an overnight culture of the bacterial strain of interest and poured over an LB-agar plate. Phage lysates were serially diluted 10-fold in LB 8 times and 2 µL of each dilution were spotted onto the top agar plates and incubated for 6-20 hours at 30°C for the host range analysis **(Fig 1D)** and 37°C for all other assays. We found that StAb3 is only able to consistently form visible plaques when using a lower percentage of top agar and incubated at lower temperatures. Therefore, StAb3 was instead plated on top agar plates prepared with 0.2% agar and 25µL of an overnight culture of the bacterial strain of interest and poured over an LB-agar plate. These plates were incubated at room temperature (∼23°C). Additionally, for both StAb2 and StAb3, using well hydrated plates by minimizing drying time for top-agar plates drastically improved our ability to consistently visualize plaques. Following incubation, plates were observed for plaques and areas of clearance. If individual plaques were observed, they were counted in the most concentrated dilution that was still countable and used to calculate PFU/mL. For some combinations of phage and bacteria, in spots where the phage was highly concentrated, the entire spot cleared showing evidence of lysis; however, as the phage was diluted, it never formed individual plaques. In the host range analysis, when this occurred, the least concentrated spot that still had clearance was noted, and as a proxy for a PFU/mL count was assumed to have the equivalent of 20 individual plaques and used to calculate PFU/mL, and when the data was reported, are marked with a black dot. For all other plaque assays, when this occurred, it was noted, PFU/mL was reported as the limit of detection (5000 PFU/mL), and they are marked with c indicating clearance. For the other strain and bacteria combinations, no clearance was observed at any dilution, PFU/mL was reported as the limit of detection (5000 PFU/mL), and they are marked with nd indicating that phage lysis was not detected. All plaque assays were performed with three independent biological replicates.

### Transmission electron microscopy

For analyses of phages at the ultrastructural level, samples were allowed to absorb onto freshly glow discharged formvar/carbon-coated copper grids (200 mesh, Ted Pella Inc., Redding, CA)) for 10 min. Grids were then washed two times in dH2O and negative stained with 1% aqueous uranyl acetate (Ted Pella Inc.) for 1 min. Excess liquid was gently wicked off and grids were allowed to air dry. Samples were viewed on a JEOL 1200EX transmission electron microscope (JEOL USA, Peabody, MA) equipped with an AMT 8 megapixel digital camera (Advanced Microscopy Techniques, Woburn, MA).

### Phage sequencing and genomic analysis

A 1 mL sample of phage lysate was treated with 10 µL (20 units) of Turbo DNase (Thermo Fischer Scientific) and 100 µL of TURBO DNase Buffer, incubating at 37°C for 30 minutes to remove any bacterial DNA. DNase was inactivated by adding EDTA to a final concentration of 15mM and heating at 75°C for 10 minutes. DNA was then purified utilizing a Phage DNA Isolation Kit (Norgen Biotek, SKU 46800). Purified DNA was prepared for Illumina sequencing utilizing Nextera Tagmentation reagents, as described previously[67]. Sequencing was performed at the Washington University Center for Genome Sciences and Systems Biology DNA Sequencing Innovation Lab following standard procedures on an Illumina MiniSeq, 2×150 paired-end, resulting in ∼50,000-150,000 reads per genome.

Demultiplexed reads were processed utilizing Trimmomatic 0.39 to remove adapter sequences, then genomes assembled with SPAdes 4.0.0 with default parameters[68,69]. Genomes were assessed using Bandage to confirm purity and circular permutation, indicating completeness of the genome sequence[70]. Genomes were manually reordered and reoriented to maximize synteny with closest relatives, as identified through BLAST searches of the complete genomes[71]. Whole-genome phylogenetic trees based on protein sequences were generated using VIPTree[72]. Tree was rendered using SplitsTree App 6.0.0; the Show Trees method was used (default options) so as to obtain a Tree View visualization[73]. Taxonomic assignment was performed using taxMyPhage web version 3.3.6[74]. Genomes were annotated using Pharokka 1.7.3 and Phold 0.2.0, with default parameters[75–79]. Clinker 0.0.31 was used to generate phage genome comparison maps[80].

Putative depolymerases of StAb2 were identified using PhageDPO Galaxy version 0.1.0 and DePP web version 1.0.0 utilizing Pharokka-annotated ORFs[50,81]. Homologues of putative depolymerases were identified in other *Lazarusvirus* and *Hadassahvirus* isolates via BLASTP. Multiple sequence alignments were then generated using Clustal Omega on the EMBL-EBI Job Dispatcher[82].

### Density gradient

A silica-based density gradient was utilized to semi-quantify capsule production using a modified version of previously published protocols[37,83]. Bacterial strains of interest were streaked onto an LB-agar plate and incubated for 16 hours at 37°C. Bacteria was scraped from the plate with a pipette tip and resuspended in PBS (Gibco, 70011-004) and normalized to an OD_600_ of 3. 1 mL of bacteria in PBS was centrifuged for 5 minutes at 5,000 x g in a 1.5mL microcentrifuge tube, and the supernatant was removed. The pellet was resuspended in 675 µL of PBS, then 325 µL of LUDOX HS-30 colloidal silica (Sigma-Aldrich, 420824) was added and mixed by inversion. The mixture was centrifuged for 30 minutes at 9000 x g. The microcentrifuge tubes were photographed and the distance from the bottom of the microcentrifuge tube to the band of bacteria was measured.

### Acquisition of capsule mutant *A. baumannii*

The plasmids and primers used in this study are listed in S7 and S8 Tables respectively. The capsule mutant for UPAB1 was generated by removing the *wzy* gene by allelic exchange as previously described[84,85]. Briefly, an approximately 300bp region upstream and downstream of *wzy* was amplified from UPAB1 genomic DNA. These regions were fused to an FRT site-flanked apramycin resistance cassette from pKD4-Apr using overlap extension PCR. This DNA was electroporated into UPAB1 carrying pAT04, a plasmid that encodes a RecAB recombinase and a hygromycin B resistance cassette followed by selection on LB-agar with apramycin. The strain was cured of pAT04 by serially passaging, isolating colonies and patching on LB-agar and LB-agar with hygromycin B until an isolate was identified that was no longer resistant to hygromycin B. The apramycin cassette was then removed by introducing the plasmid pAT03, which encodes an IPTG-inducible FLP recombinase, then plating on LB-agar with hygromycin B and 2mm IPTG. Successful removal was verified by confirming a lack of growth on LB-agar with apramycin and PCR amplification of the deleted region. The strain was cured of the pAT03 plasmid by passaging, plating for individual colonies, and patching on LB and hygromycin B to identify an isolate which is no longer resistant to hygromycin B.

The capsule mutants of Ab5075, AbCAN2, and Ab014 were acquired spontaneously. Upon streaking freezer stocks of these strains on an LB-agar plate for isolation of individual colonies, two morphologies were observed, a primary morphology that is more opaque and mucoid and another, less frequent morphology that is more translucent and non-mucoid. It was predicted that the mucoid colonies were the wild-type capsulated strains while the non-mucoid colonies had lost their capsule. These two colony types were isolated and assessed for capsule production by the density gradient described above. All of the capsule mutants were further validated by sequencing as described below. K-locus type for each wild-type strain was identified using Pathogenwatch[24,86].

### Generation of UPAB1 Δ*wzy* transposon mutagenesis library

A transposon mutant library in UPAB1 Δ*wzy* was prepared using a modified version of vector pJNW684 which encodes the Himar1 mariner transposon system as described previously[87,88]. Plasmids and primers used in this study are listed in S7 and S8 Tables respectively. The selection marker of pJNW684 is originally a kanamycin resistance cassette; however, many *A. baumannii* strains, including UPAB1 are resistant to this antibiotic; therefore, it was exchanged for an apramycin resistance cassette. The apramycin resistance cassette was amplified from pUCT18T-miniTn7T-Apr, then introduced into pJNW684 using NEBuilder HiFi DNA Assembly Master Mix (New England Biolabs). The transposon was introduced into UPAB1 Δ*wzy* through conjugation from *E. coli* S17-1 λ-pir carrying pJNW684-Apr. Conjugation was facilitated by the conjugal helper plasmid pRK2013 carried in *E. coli* HB101 and transposon insertion was facilitated by the plasmid pTNS2 carried by *E. coli* 100D. UPAB1 Δ*wzy* and each *E. coli* strain were grown overnight in LB at 37°C in the appropriate antibiotic. 35 mL of each strain was centrifuged at 6500 rpm for 3 minutes and washed once with LB, centrifugating at 6500 rpm for 3 minutes. The pellets of all bacteria were resuspended in 1.4 mL of LB and combined, and 20uL of this bacteria mixture was spotted 70 times on LB-agar plates for conjugation to occur. Plates were dried for 30 minutes at room temperature them incubated at 37°C for 1 hour. To select for UPAB1 Δ*wzy* isolates with transposons inserted, the bacterial spots were resuspended in LB and spread on LB-agar plates with 10 ug/mL chloramphenicol and 50 ug/mL apramycin and incubated overnight at 37°C.

### Selection of StAb2-resistant UPAB1 Δ*wzy* transposon mutants

In order to identify bacterial genes essential for infection by StAb2, UPAB1 Δ*wzy* transposon mutants from the library generated above that are resistant to StAb2 were isolated. 100 mL LB with 60 ug/mL apramycin was inoculated with 100 uL (4,000 CFU) of the transposon mutagenesis library and grown at 37C shaking, after 1, 3, and 8 hours, 3.5×10^8^ PFU of StAb2 was added, and after 3, 8, and 24 hours, a sample of the culture was taken and plated on LB-agar plates with 60ug/mL apramycin and 10ug/mL chloramphenicol to obtain isolated colonies. Five isolates from each timepoint were tested for susceptibility to StAb2 by plaque assay. All isolates were resistant to StAb2. Genomic DNA was purified from one isolate from the 3-hour timepoint, and 2 isolates each from the 8 and 24 hour timepoints using the PureLink Genomic DNA Mini Kit (Thermo Fisher Scientific). DNA was sequenced by Illumina sequencing by SeqCenter and the location of transposon insertion was determined using Geneious Prime 2024.0.7 (https://geneious.com).

### Bacterial growth curves

To assess the influence of treatment with bacteriophages on bacterial growth, growth curves were performed. Solutions containing overnight culture of the indicated bacterium normalized to an OD_600_ of 0.01 and bacteriophage lysate was added to the indicated MOIs (with an MOI of 1 corresponding to 2×10^6^ PFU/mL). When multiple bacteriophages were used in a single treatment, they were added at equal MOIs to have a total of the reported MOI. For example, if two phages were added at a total MOI of 1, then an MOI of 0.5 for each phage was added. 150 µL aliquots of these solutions were added to the wells of a polystyrene 96-well plate (Corning, 3788). Plates were incubated at 37°C with shaking and OD_600_ was measured every 30 minutes in a microplate reader (Accuris Smartreader 96-T or BioTek Synergy HTX). Each combination of phages and bacteria was performed with three independent biological replicates, each graphed independently. The growth curves in 7C, 7D, and S5A-B were performed using the Accuris Smartreader 96-T (MR9600-T) each with 6 technical replicates shaking at 4.7 Hz. The growth curves in 7E and S5D Replicate 2 were performed with the same settings but with 3 technical replicates. The growth curves in S5C and S5D Replicate 3 were performed using the BioTek Synergy HTX with orbital shaking at 236 cpm and an amplitude of 4mm with 3 technical replicates.

### Generation of escape mutants

398 mutants resistant to StAb1 (398StAb1eA-H) or StAb2 (398StAb2eB-H), 17978 Δ*pglC* mutants resistant to StAb2(17978 ΔpglC-StAb2E1-2), and G7 mutants resistant to Bhz16 were acquired as follows. 150 µL aliquots of LB with an overnight culture of the indicated bacterium normalized to an OD_600_ of 0.01 and the bacteriophage of interest were incubated at 37°C in a 96-well plate (Corning, 3788) shaking for 16-24 hours until visibly turbid. Bacteria from the wells were streaked onto an LB-agar plate to obtain isolated colonies. An individual colony was isolated and durable resistance to the bacteriophage of interested was assessed with a plaque assay as described above. When MC47.2 was grown with StAb3 in this manner, bacterial growth occurred; however, when isolates were assessed, they remained susceptible to StAb3. To acquire MC47.2 mutants resistant to StAb3, 4 mL of top agar (LB with 0.4% agar) was combined with 25 µL of overnight culture of the indicated bacterium and poured over LB-agar plates and LB-agar plates supplemented with chloramphenicol. A 10µL drop of StAb3 phage lysate was spotted onto the top agar plates and incubated at 30°C or 20°C until individual colonies appeared in the area of clearance. Colonies were selected and streaked onto a fresh LB-agar plate to isolate individual colonies, and resistance to StAb3 was assessed by plaque assay. The Up280 mutant resistant to StAb3 was acquired in a similar manner; however, 0.2% agar was used for the top-agar instead of 0.4% and plates were incubated at room temperature (∼23°C). To acquire one of the 398 mutants resistant to StAb2 (398-StAb2eA), 4 mL of top agar was combined with 25 µL of overnight culture of 398 and poured over LB-agar plates. StAb2 was spotted on this plate, and then plates were incubated at 37°C until colonies appeared in the area of clearance. Colonies were selected and streaked onto a fresh LB-agar plate to isolate individual colonies and resistance to StAb3 was assessed by plaque assay.

### Sequencing and analysis of capsule mutants and phage resistant strains

Genomic DNA was purified from each capsule mutant, escape mutant, and relevant wild-type strains utilizing the PureLink Genomic DNA Mini Kit (Thermo Fisher Scientific). DNA was sequenced by Illumina sequencing by SeqCenter or SeqCoast. The Illumina sequencing reads for sequenced strains were mapped to the assembled genome listed with Geneious Prime 2024.0.7 (https://geneious.com) utilizing the Map to Reference function with the Geneious Mappper at Medium-Low Sensitivity/Fast. Following assembly, the Find High/Low Coverage function was used to find regions with coverage below 4 standard deviations from the mean and the Find variations/SNPs function was used to identify variants with minimum variant frequency set to 0.2, maximum variant P value set to 10^-6^, and minimum strand-bias P-value set to 10^-5^ when exceeding 65% bias finding variants “Inside & Outside CDS” with the “Approximate P-value” calculation method. SNPs identified with variant frequency >70% were reported as a SNP. Areas of the genome with SNPs of variant frequency <70% or coverage below 4 standard deviations from the mean were manually assessed for evidence of rearrangements, insertions, or deletions. When an assembled genome for the wild-type strain or a closely related strain was available, Illumina sequencing from mutants and their associated wild-type strain was aligned to this genome, and the results from the wild-type strain were used to eliminate any spurious calls. For 398, MC47, and Up280, which did not have an assembled genome, SeqCenter’s Small Nanopore Combo service was used to perform Nanopore and Illumina sequencing and acquire an assembled genome to which all escape mutants from these strains were aligned. For G7, DNA was sent to Plasmidsaurus for Nanopore sequencing and genome assembly, to which all escape mutants were aligned, and Illumina sequencing for the wild-type strain was used to eliminate any spurious calls. For Ab014, sequencing was performed by McGill University and the Genome Quebec Innovation Center with PacBio, to which Illumina sequences of Ab014 and Ab014-cm were aligned and compared. K-locus annotations to characterize the SNPs found in 398 and G7 escape mutants were performed using Kaptive[24].

### Strain construction

Plasmids and primers used in this study are listed in S7 and S8 Tables respectively. The deletion of *carO* from 398 and 398-StAb1eA was generated using a pEX-based exchange vector with counterselection as described previously[89,90]. In brief, regions approximately ∼1kb upstream and ∼1kb downstream of *carO* were amplified from the 398 genome and were cloned into pEX18-Apr with NEBuilder HiFi DNA Assembly Master Mix (New England Biolabs). The plasmids were introduced into 398 and 398-StAb1eA using a tri-parental mating with *E. coli* Stellar Competent Cells carrying pEX18Ap_carO398 serving as the donor, and *E. coli* HB101/pRK2013 facilitating conjugation. Acinetobacter with successful integration of the plasmid were selected for on LB-agar supplemented with apramycin and chloramphenicol. Integration of the plasmid into the expected location was validated by PCR, then strains were plated on LB-agar with no NaCl and 10% sucrose to select for excision of the plasmid. Isolates that had the desired loss of *carO* were identified utilizing PCR with primers flanking *carO*.

Plasmids and primers used in this study are listed in S7 and S8 Tables respectively. Complementation of *carO* in UPAB1 Δ*wzy-*StAb2TnR1 was conducted by inserting the *carO* gene from four different *A. baumannii* strains into a neutral site in the chromosome with a miniTn7 system as previously described[91,92]. The constructs were generated by amplifying the putative promoter region (∼500bp upstream) along with the open reading frame of the *carO* genes from 19606, AYE, 398, and ACICU, which was then inserted into pUCT18T-mini-Tn7T-Zeo with NEBuilder HiFi DNA Assembly Master Mix (New England Biolabs) following cleavage of the plasmid with KpnI and BamHI. The plasmids were introduced into UPAB1 Δ*wzy-*StAb2TnR1 using a four-parental conjugation technique and complemented strains were selected for on LB-agar supplemented with zeocin and chloramphenicol, and insertion was validated by PCR[91,92].

### Adsorption assays

Cultures of strains to be assessed for phage adsorption were inoculated from streaks on LB agar plates into LB and cultured overnight at 37°C. The following morning, the indicator strain on which plaquing is performed (Ab014Δcps) was inoculated into LB and cultured until mid-late log phase was reached (OD∼1; 2-3 hours). 25 μL of this culture was used to seed 4 mL of LB top agar (0.4% agar) per agar plate and then allowed to solidify. Simultaneously, overnight cultures of assayed strains were diluted 1:100 into 5 mL of LB broth in 125-mL flasks and incubated with shaking at 37°C until early-mid log was reached (OD ∼0.3-0.6; corresponding to ∼5×108 CFU/mL of bacteria). Flasks containing no bacteria, only LB, were treated similarly as a negative control to identify any background carryover of phage. 10^6^ PFU/mL of phage StAb2 was then added to the flask and allowed to incubate with shaking at 37°C for 10 minutes. Flasks were immediately transferred onto ice to rapidly cool, and samples were kept on ice for the remainder of processing. 500 μL aliquots were transferred to 1.5 mL microcentrifuge tubes and centrifuged to pellet bacteria (17,000 x g, 2 minutes, 4°C). 100 μL aliquots were removed (supernatant), and the pellet was washed in 500 μL fresh, cold LB and pelleted again. The supernatant was removed and the pellet resuspended in 500 μL fresh, cold LB. 100 μL aliquots were removed (pellet). Supernatant and pellet were both 10-fold serially diluted 5 times, and 20 μL spots of each dilution were plated onto the top agar overlay of the indicator strain. These plates were incubated overnight at 30°C within a BD GasPak EZ containing a damp paper towel to maintain sufficient moisture for consistent StAb2 plaquing. Plaques were counted and used to determine percent binding of phage: (PFU/mL of pellet / (PFU/mL of pellet + PFU/mL of supernatant)) * 100.

### NGR-like exopolysaccharide locus analysis

Genes in the predicted NGR locus in *A. baumannii* were compared to genes from other exopolysaccharide synthesis and transport systems. Percent identity between proteins listed in Figure S3 was determined using UniProt align, which makes use of Clustal Omega on the EMBL-EBI Job Dispatcher[82,93–96]. Structural alignment proteins with existing structure predictions through AlphaFold shown in S4 Table was performed using RCSB PDB pairwise structure alignment with the TM-align method[97].

## Acknowledgements

We thank the Washington University Center for Genome Sciences and Systems Biology DNA Sequencing Innovation Lab for help with sequencing. We thank Wandy Beatty for help with TEM imaging of phages. We thank Dr. Clay Jackson-Litteken and Dr. Manon Janet-Maitre for generating plasmid pJNW684-Apr. We thank members of the Feldman and LeRoux labs for helpful discussions.

## Supporting information

**S1 Fig.**
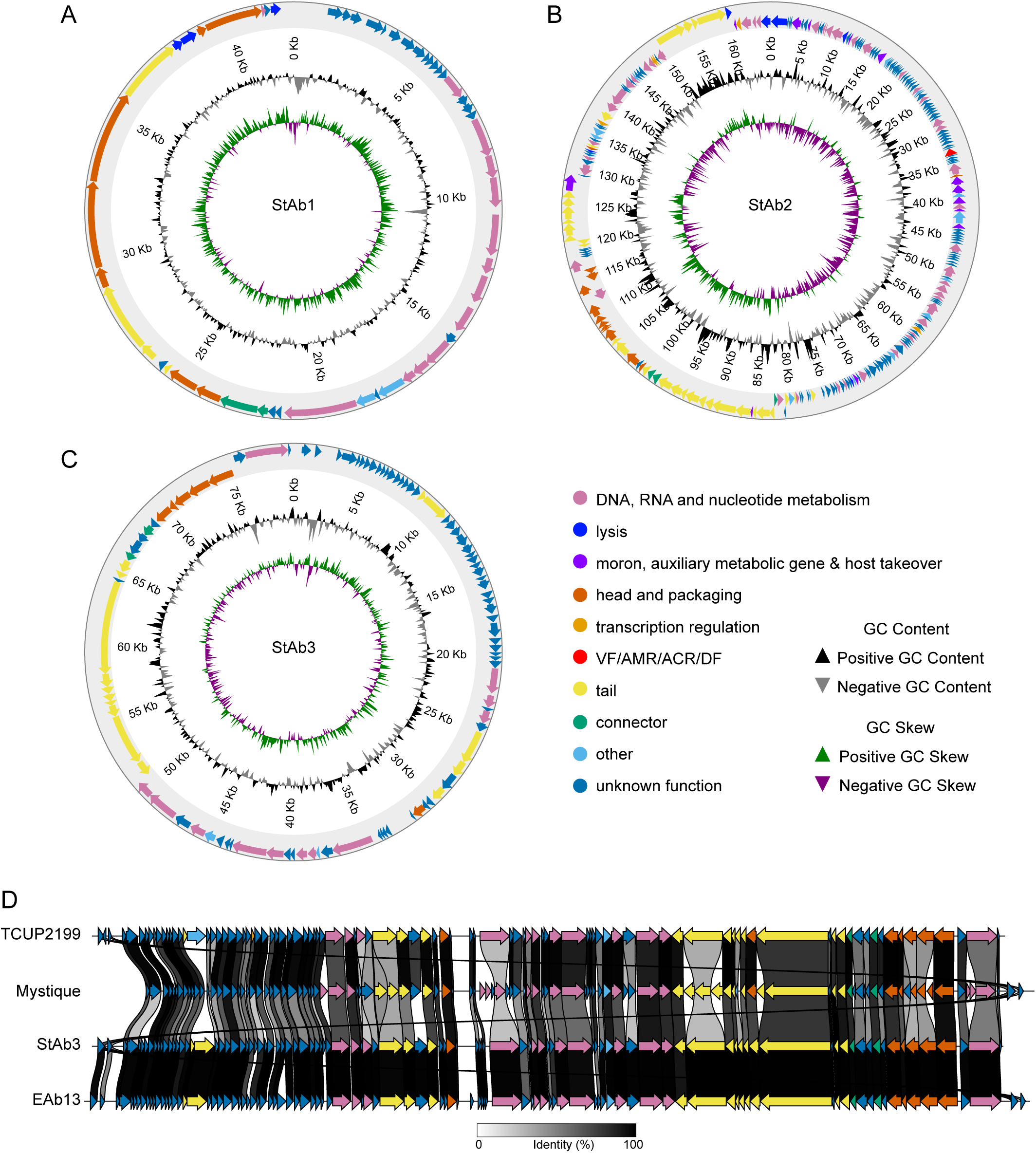
Genomic characteristics of novel *Acinetobacter* phages StAb1, StAb2, and StAb3. A-C) Circularly permutated diagrams of the genomes of StAb1, StAb2, and StAb3, generated with Phold. ORFs are annotated and colored according to PHROG predictions as indicated. Interior tracks indicate the GC content and skew across the genomes. D) Comparison of annotated ORFs in StAb3 with its closest sequenced relatives: TCUP2199, Mystique, and EAb13, generated using Clinker. Phold-annotated ORFs are identified and colored as in (A). Protein percent identity is indicated with the black lines between each gene and its homolog in the adjacent tracks.

**S2 Fig.**
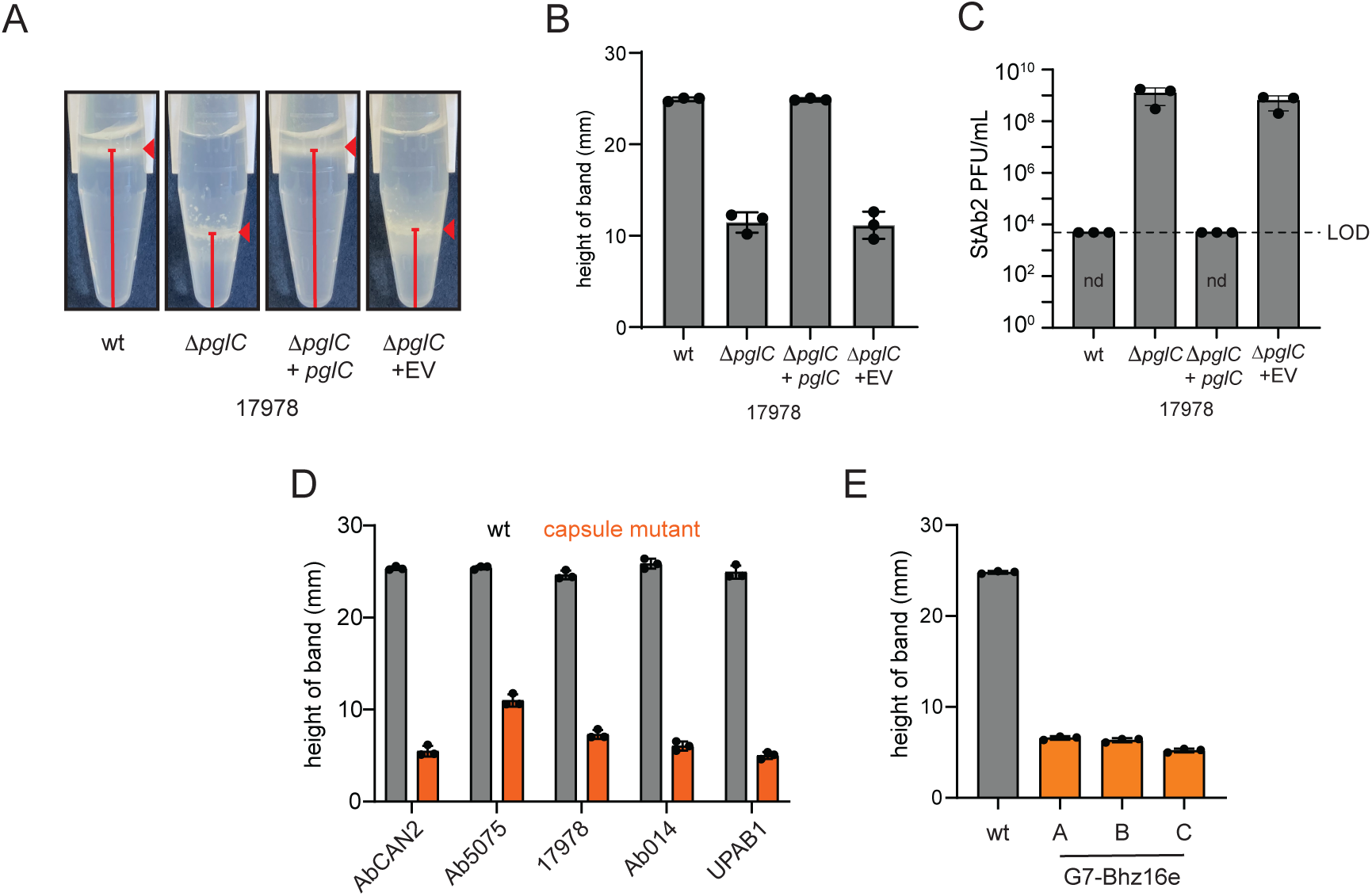
StAb2 can infect acapsular strains. A) Density gradient assay for 17978, 17978Δ*pglC*, the complemented strain 17978Δ*pglC* +*pglC* (PRLM2), and the empty vector control 17978Δ*pglC* + EV(PWH1266). The area of bacterial biomass is indicated by the red triangle. The height of this band in mm from the bottom of the tube to the center of the bacterial biomass was used as an indicator of bacterial density and therefore capsular content as marked with a red line. B) Quantification of band height for strains of 17978 as in (A). C) Quantification of plaque assay with StAb2 on strains from (A) as measured by PFU/mL in a plaque assay. LOD = 5 x 10^3^. nd = not detected. D) Wild-type strains and corresponding capsule mutants assessed for capsule via silica-based density gradient experiments. E) Bhz16 phage escape mutant capsule formation assessed via silica-based density gradient experiments and compared with the wild-type (wt) parental strain G7 (gray). B, C-F) Three independent replicates and their average is presented with error bars representing sd.

**S3 Fig.**
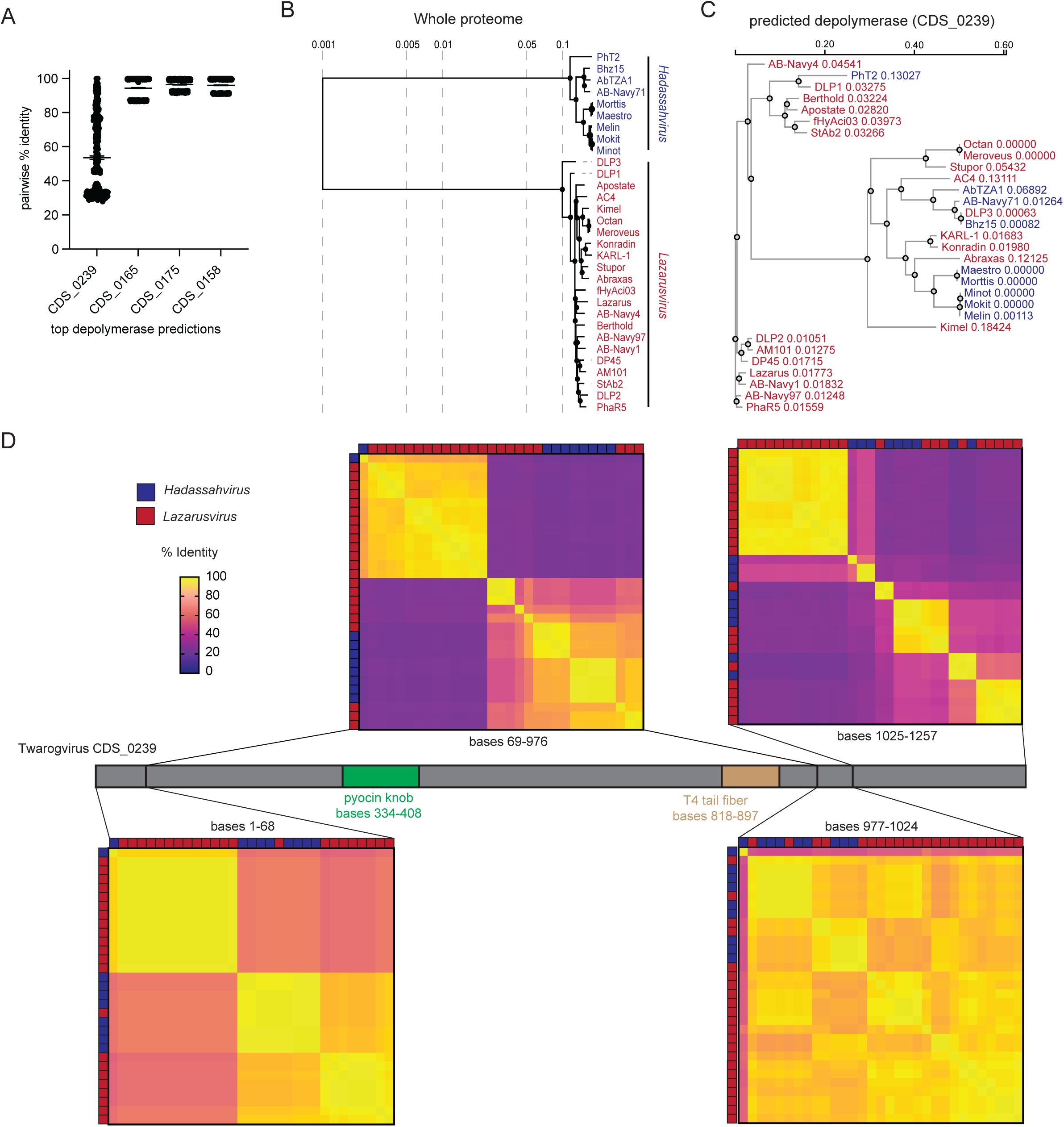
Twarogviruses carry a highly variable predicted depolymerase that may explain host range restriction. A) The pairwise percent identity comparisons of the top four most strongly predicted depolymerase genes in StAb2 compared to each homolog in the 29 *Hadassahvirus* and *Lazarusvirus* genomes available on NCBI, as well as Bhz15. The protein sequence homolog was compared to every other homolog in the 31 strains using Clustal Omega, and the percent identity of each comparison is shown. B) A whole-proteome phylogenetic tree of all 29 *Hadassahvirus* and *Lazarusvirus* genomes available on NCBI, as well as StAb2 and Bhz15, generated using VIPTree. C) Protein sequence-based phylogenetic trees of strongly predicted depolymerase CDS_0239 from StAb2 and its homologs. D) A diagram of CDS_0239 from StAb2, with functional domains annotated. A comparison of the protein sequence among homologs from Twarogviruses is displayed, revealing two short linker domains (aa 1-68 and aa 977-1024), with two larger highly variable domains (aa 69-976 and 1025-1257). The pairwise percent identities from Clustal Omega protein alignments are shown via heatmaps for each domain. Taxonomy of each source genome is indicated along the edge of the heatmap.

**S4 Fig.**
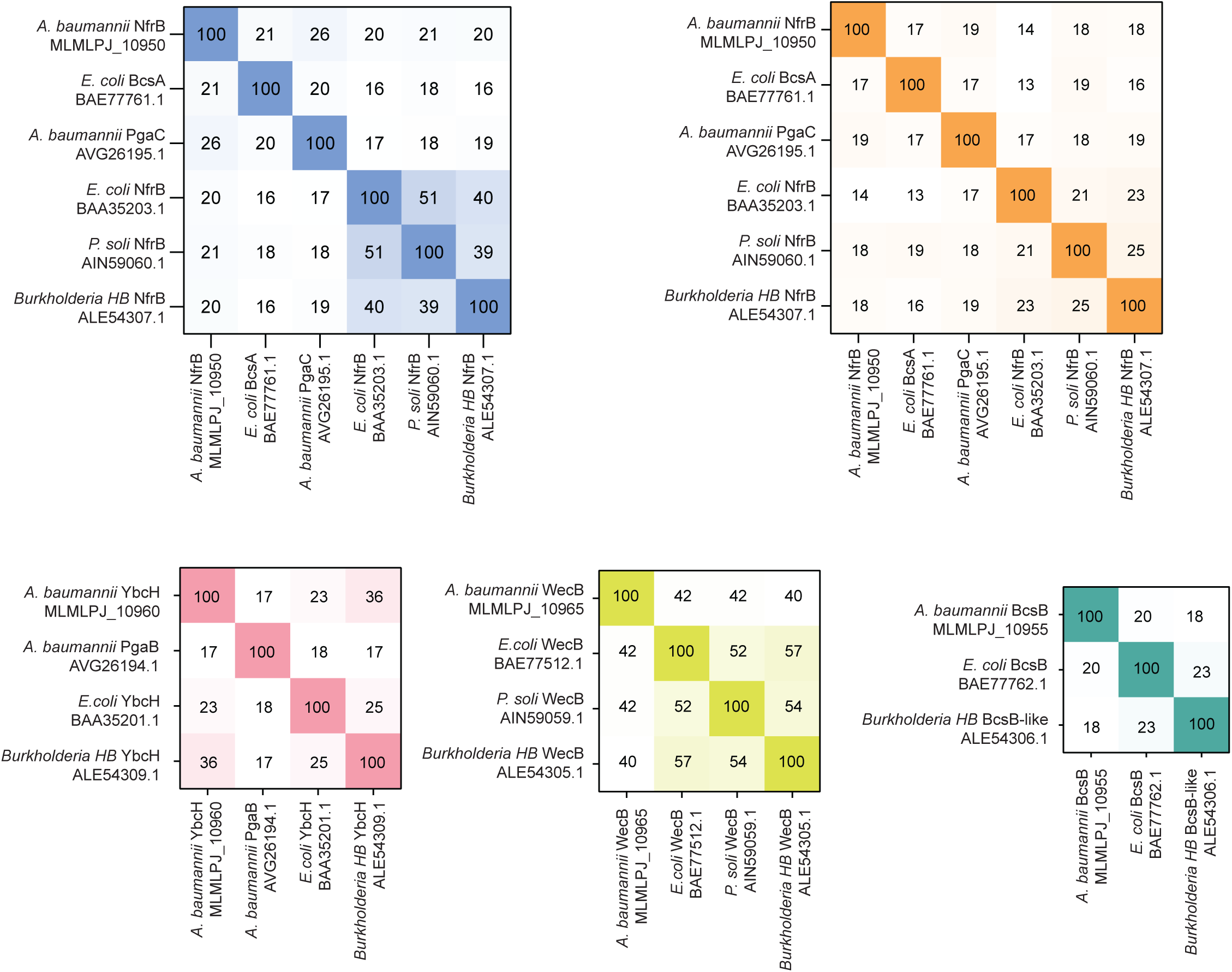
Homology between proteins involved in EPS biosynthesis and secretion pathways. The pairwise percent identity comparisons of the proteins identified as having homology in Figure 5

**S5 Fig.**
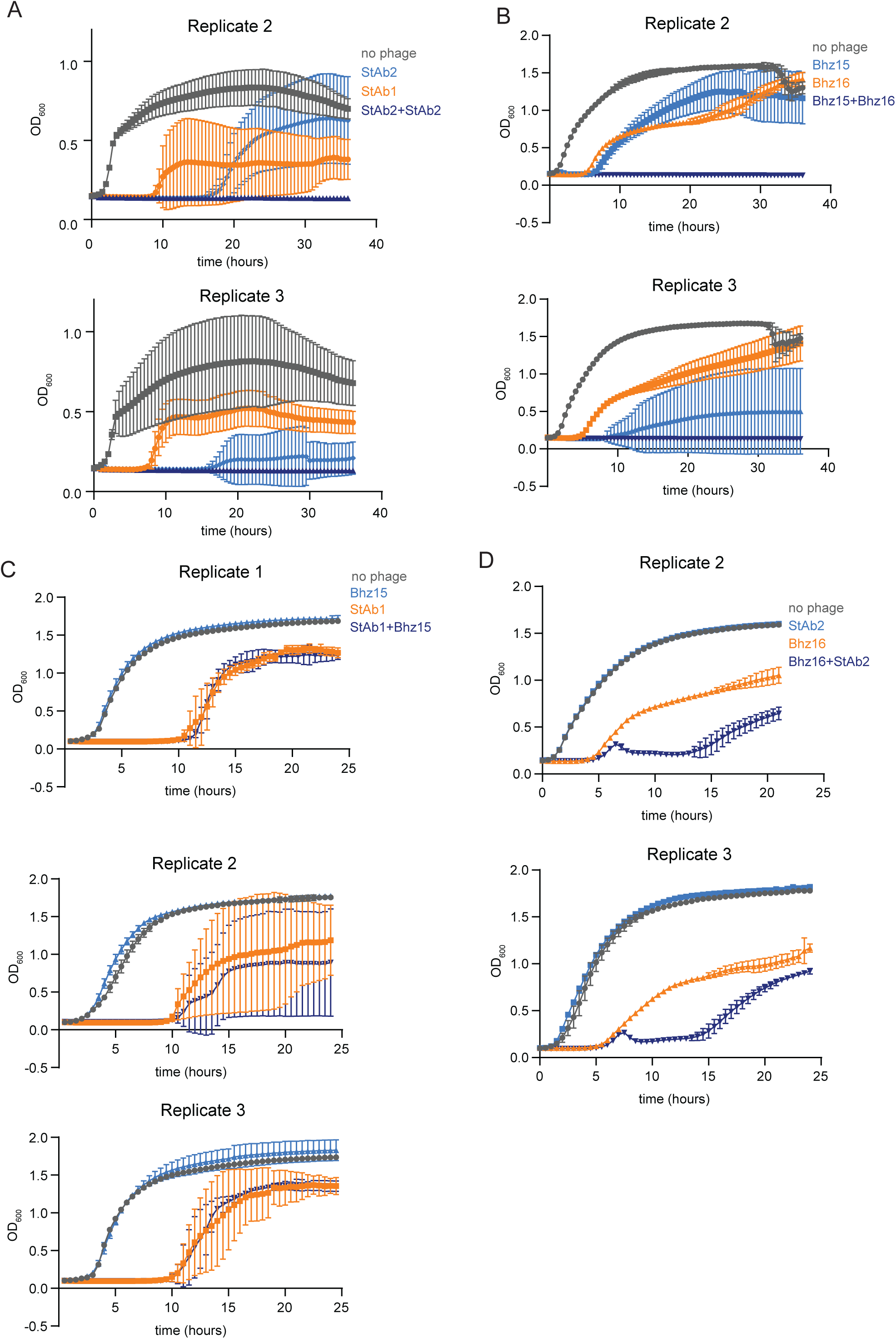
Combinations of phages prevent or delay bacterial resistance. Growth of 398 (A&C) or G7 (B&D) with the indicated individual phages or phages combined. Phages were added to an MOI of 1. Each graph represents a single independent replicate with the average of three (C&D) or six (A&B) technical replicates presented with error bars representing sd.

**S1 Table.**
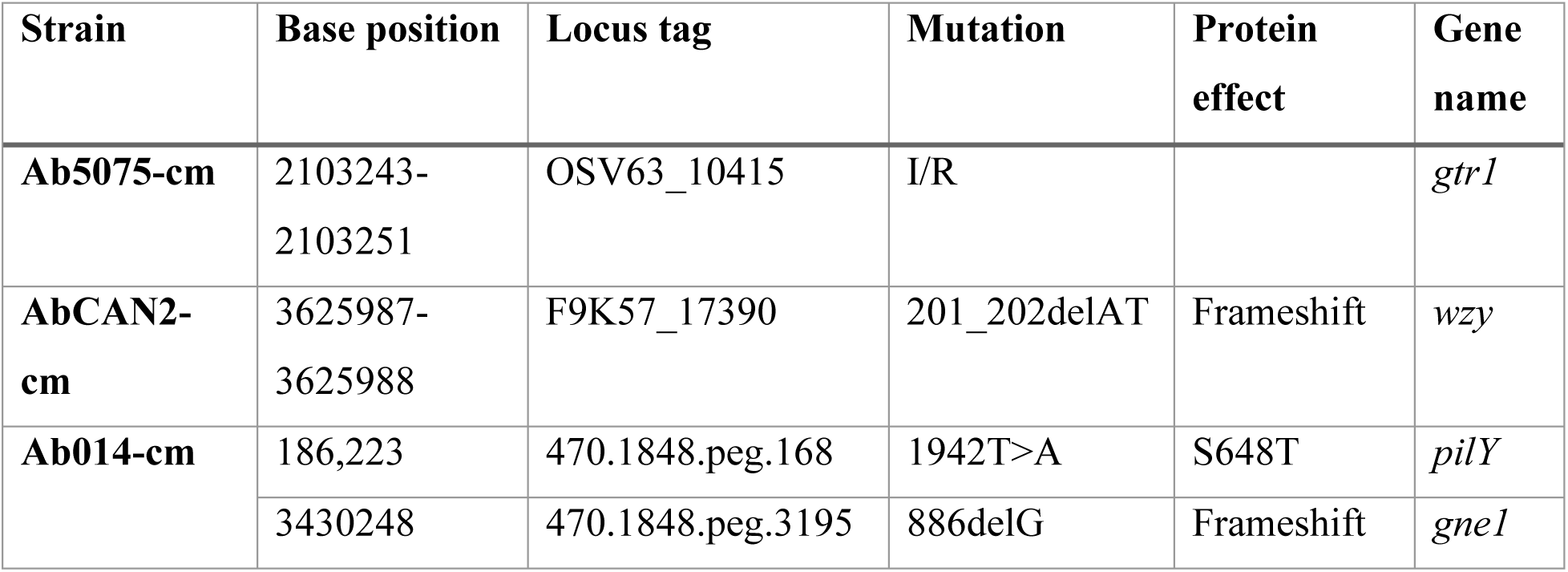
Capsule mutant confirmation sequencing analysis. Location and type of SNPs identified in spontaneous capsule mutants. Predicted insertions or rearrangements are reported as I/R.

**S2 Table.**
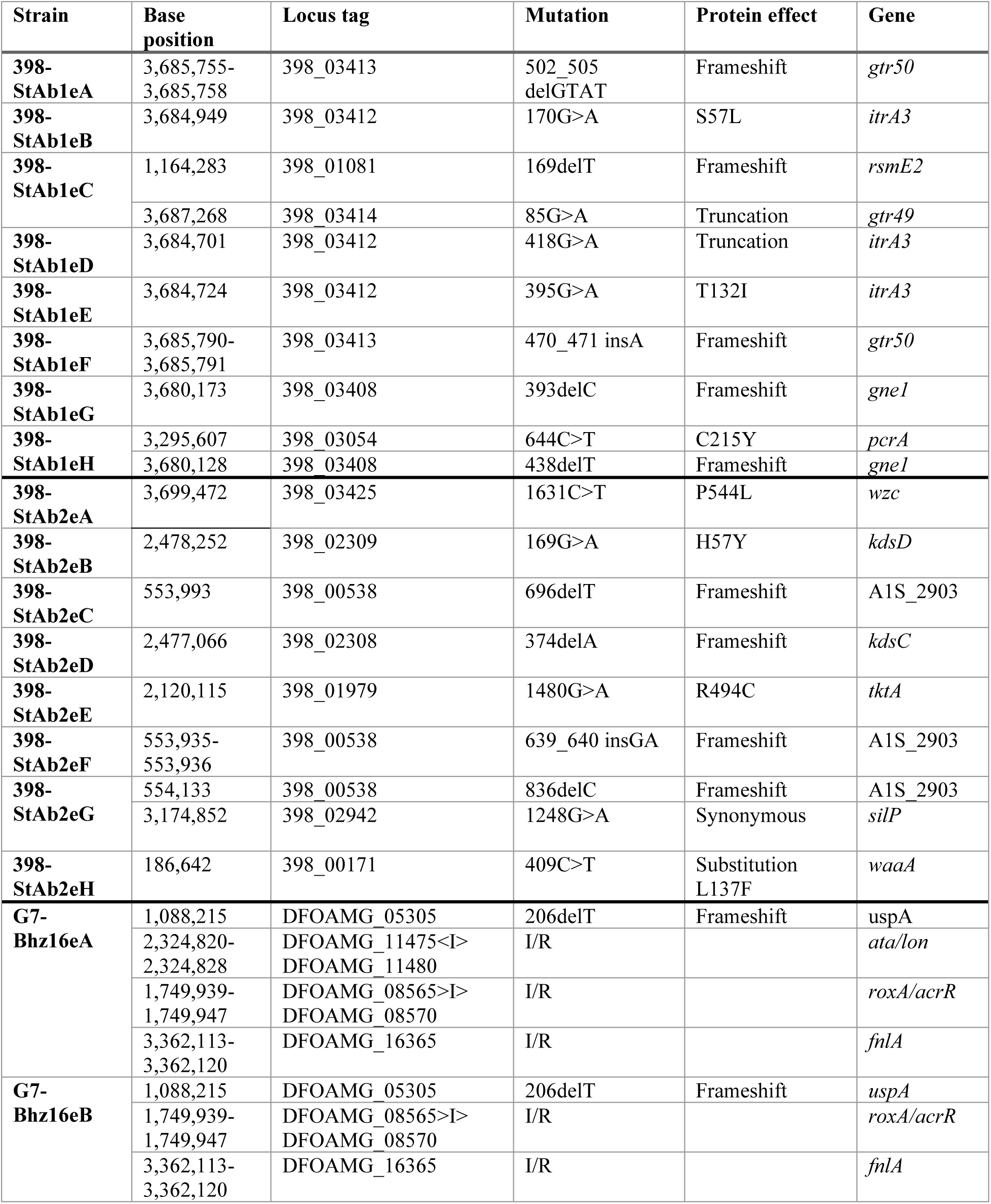
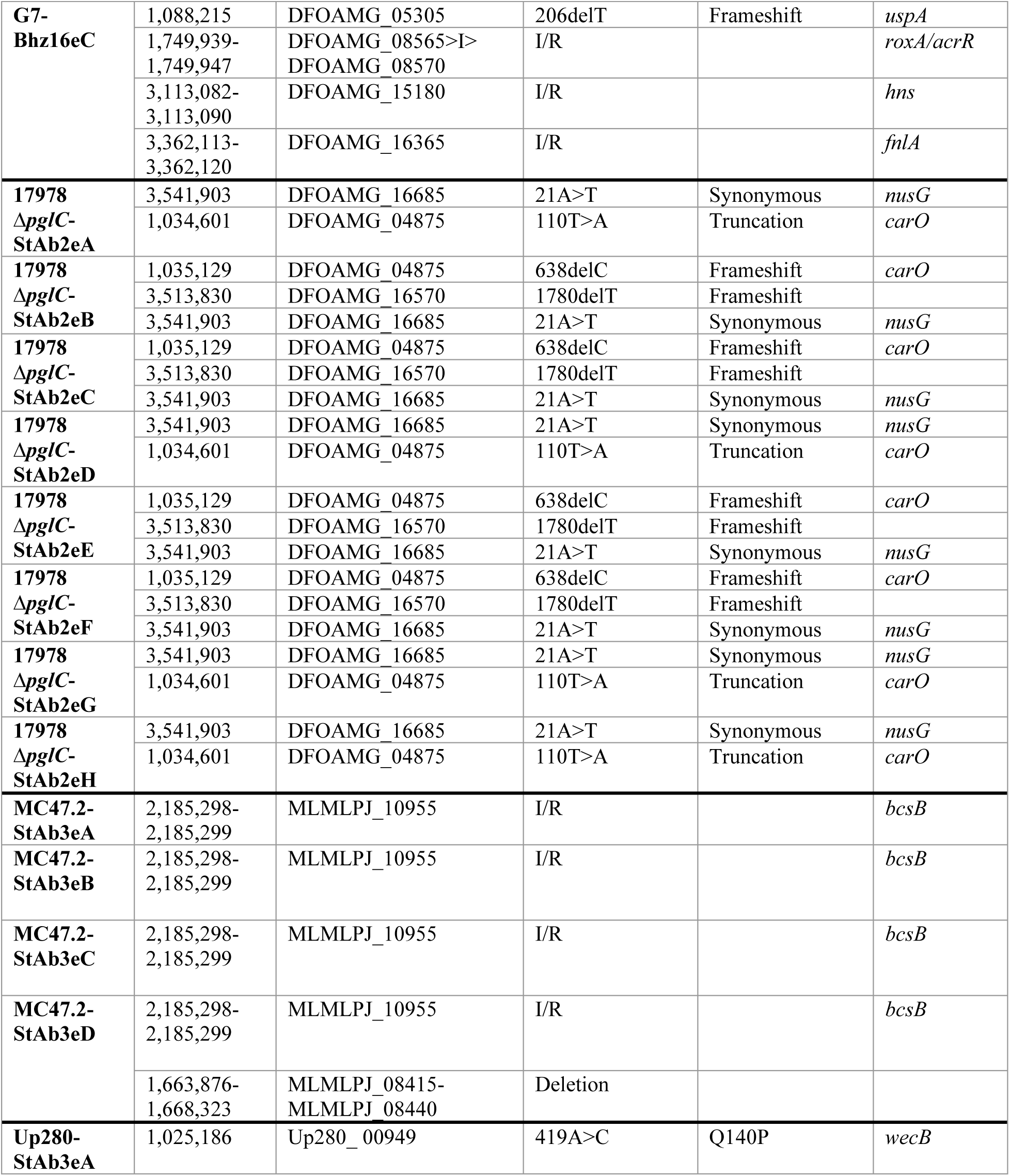
Phage-resistant mutant sequencing analysis. Location and type of SNPs identified in phage-resistant mutants. Predicted insertions or rearrangements are reported as I/R.

**S3 Table.** Capsule types of wild-type *A. baumannii* strains. K-locus and capsular polysaccharide types of *A. baumannii* strains determined using Kaptive[24].

| Strain | K-locus type | Capsular polysaccharide (CPS) type |
| --- | --- | --- |
| **398** | KL24 | K24-Wzy-GI2 |
| **MC47.2** | KL9 | K9 |
| **G7** | KL9 | K9 |
| **Ab5075** | KL25 | K25 |
| **17978** | KL3 | K3 |
| **UPAB1** | KL149 | unknown (likely K9[1,2]) |
| **AbCAN2** | KL140 | unknown |
| **Ab014** | KL120 | unknown |

---

**S4 Table.** Structural alignment of exopolysaccharide synthesis and transport proteins. Structural alignment of a subset of exopolysaccharide synthesis and transport proteins described in Figure 5B using RCSB PDB pairwise structure alignment[97].

| Aligned to | Protein | ID | RMSD | TM-Score | Identity | Aligned residues | Sequence length | Modeled residues |
| --- | --- | --- | --- | --- | --- | --- | --- | --- |
| ***A. baumannii* NfrB D0CBT6** | *E. coli* BcsA | P37653 | 3.09 | 0.82 | 17% | 347 | 872 | 872 |
|  | *A. baumannii* PgaC | C8YYH7 | 2.78 | 0.83 | 23% | 348 | 392 | 392 |
|  | *E. coli* NfrB | P0AFA5 | 3.19 | 0.86 | 17% | 363 | 745 | 745 |
|  | *P. soli* NfrB | A0A1H9LH73 | 3.22 | 0.86 | 16% | 359 | 724 | 724 |
| ***A. baumannii* BcsB D0CBT5** | *E. coli* BcsB | P37652 | 4.24 | 0.75 | 9% | 446 | 779 | 779 |
| ***A. baumannii* YbcH D0CBT4** | *A. baumannii* PgaB | C8YYH6 | 4.05 | 0.63 | 13% | 197 | 510 | 510 |
|  | *E. coli* YbcH | P37325 | 3.28 | 0.65 | 21% | 199 | 296 | 296 |
| ***A. baumannii* WecB D0CBT3** | *E. coli* WecB | P27828 | 1.64 | 0.94 | 42% | 371 | 376 | 376 |
| ***A. baumannii* NfrA D0CBT2** | *E. coli* BcsC | P37650 | 6.05 | 0.53 | 8% | 157 | 1157 | 1157 |
|  | *A. baumannii* PgaA | C8YYH5 | 6.34 | 0.48 | 6% | 155 | 747 | 747 |
|  | *E. coli* NfrA | P31600 | 4.25 | 0.67 | 11% | 258 | 990 | 990 |

**S5 Table.**
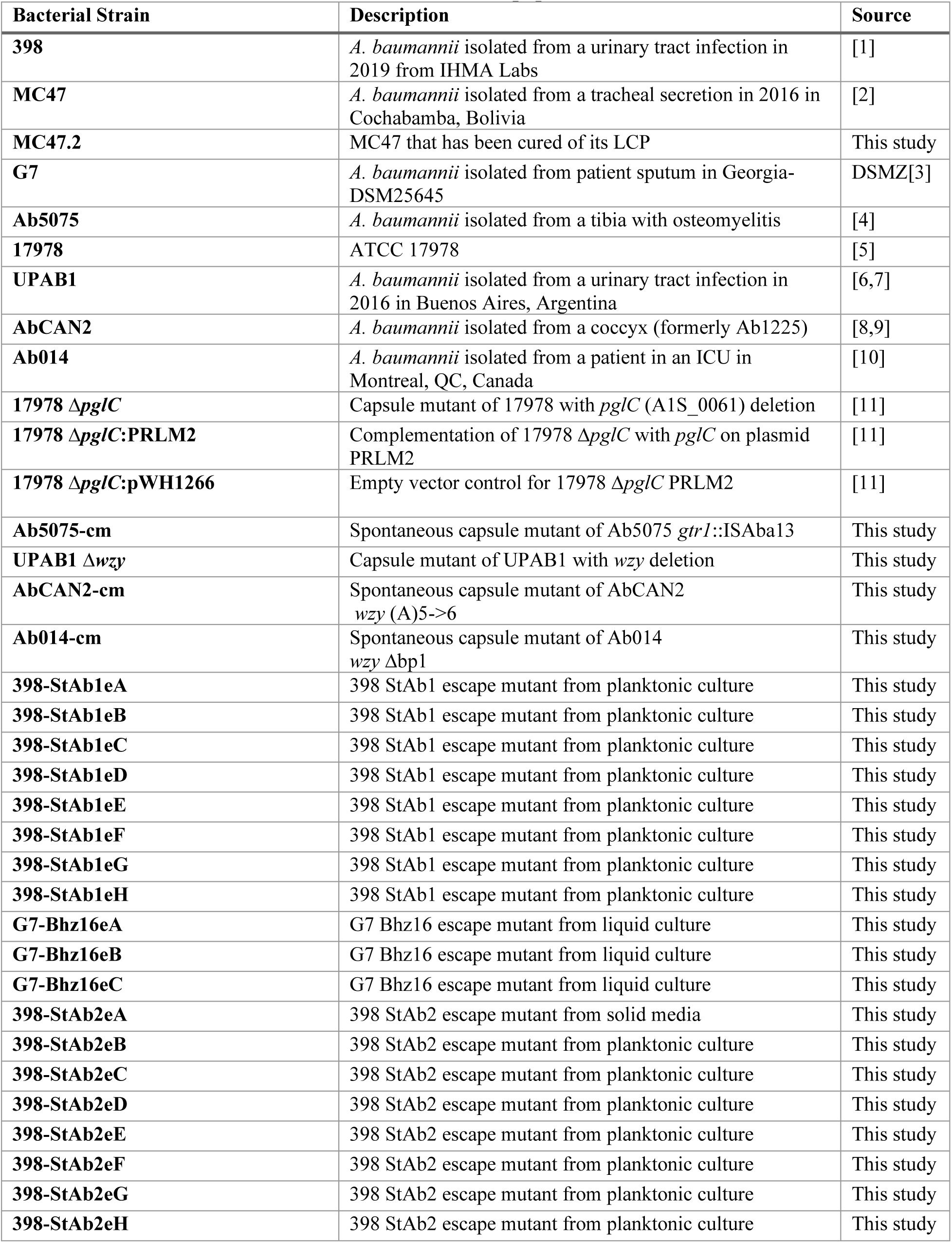
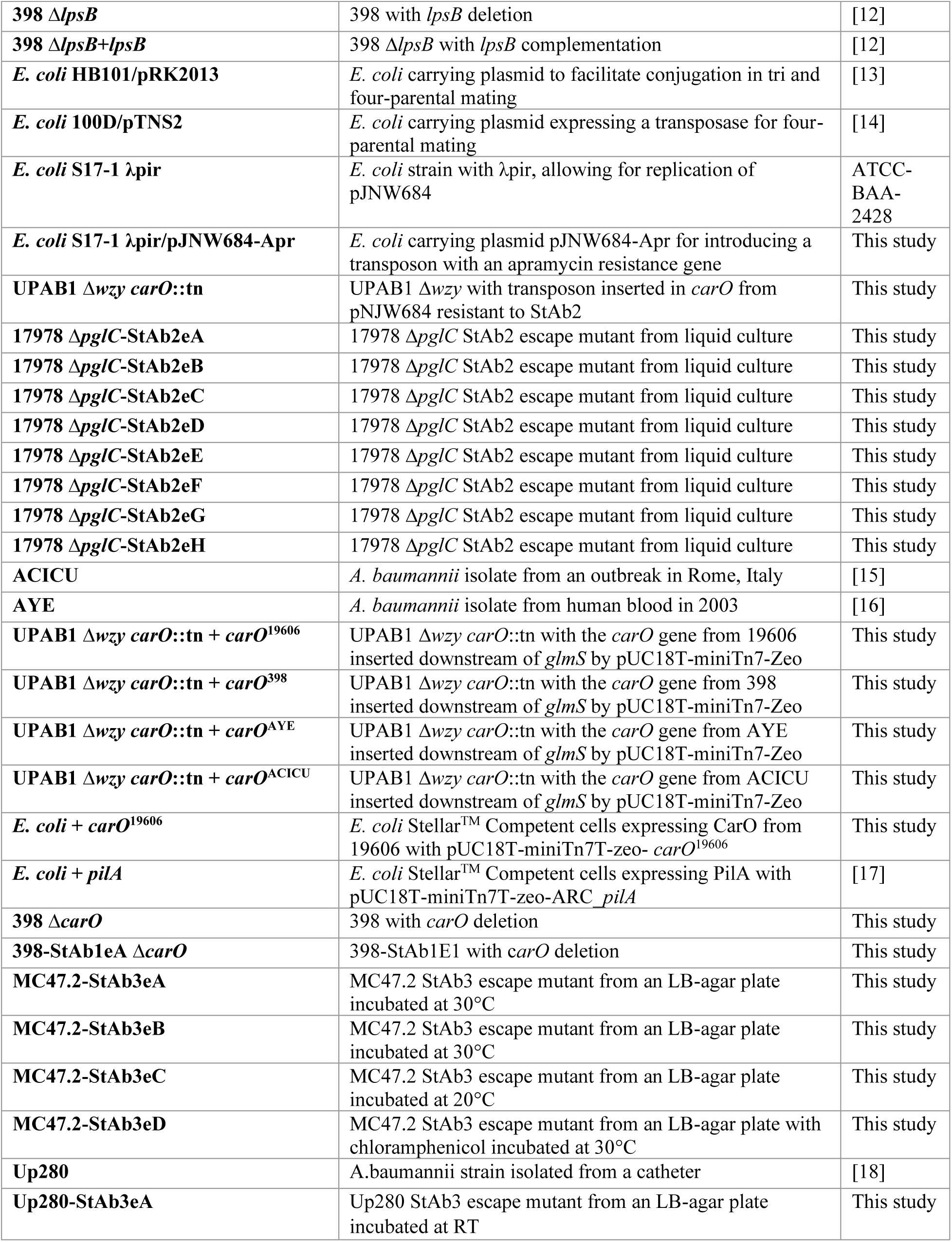
Bacterial Strains. List of all bacterial strains used and described in the paper.

**S6 Table.**
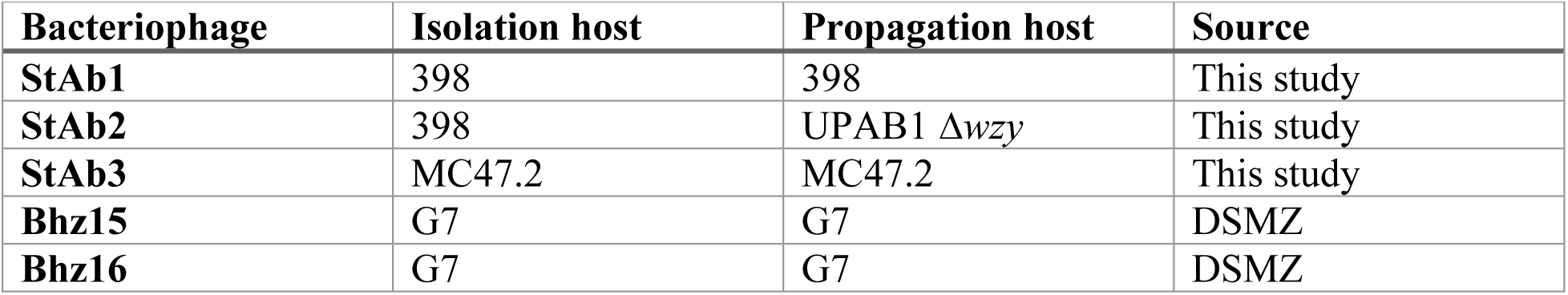
Bacteriophages. List of all bacteriophages with their isolation and propagation hosts used in this paper.

**S7 Table.** Plasmids. List of all plasmids used in the paper.

| **Plasmid** | **Description** | **Source** |
| --- | --- | --- |
| **pKD4-Apr** | Source of FRT-flanked apramycin resistance cassette for generation of UPAB1 $\Delta wzy$ | [1] |
| **pAT04-Hygro** | Plasmid encoding RecAB recombinase for facilitating allelic exchange in generation of UPAB1 $\Delta wzy$ | [1–3] |
| **pAT03-Hygro** | Plasmid encoding IPTG-inducible copy of FLP recombinase to remove the antibiotic resistance cassette for generation of UPAB1 $\Delta wzy$ | [1–3] |
| **pJNW684** | Plasmid encoding the Himar1 mariner transposon system to generate transposon insertion mutants in *A. baumannii* | [4] |
| **pUC18T-miniTn7T-Apr** | Source for apramycin cassette for pJNW684-Apr | [5] |
| **pJNW684-Apr** | pJNW684 with the kanamycin resistance cassette exchanged for an apramycin resistance cassette | This study |
| **pUC18T-miniTn7T-LAC-zeo** | Vector carrying a mini-Tn7 system for insertion of genes in the *A. baumannii* chromosome downstream of the glmS gene | [6] |
| **pUC18T-miniTn7T-zeo-ACICU_carO** | pUC18T-miniTn7T-zeo with *carO* amplified from ACICU for complementation | This study |
| **pUC18T-miniTn7T-zeo-AYE_carO** | pUC18T-miniTn7T-zeo with the *carO* amplified from AYE for complementation | This study |
| **pUC18T-miniTn7T-zeo-19606_carO** | pUC18T-miniTn7T-zeo with the *carO* amplified from 19606 for complementation | This study |
| **pUC18T-miniTn7T-zeo-398_carO** | pUC18T-miniTn7T-zeo with the *carO* amplified from 398 for complementation | This study |
| **pUC18T-miniTn7T-zeo-ARC_pilA** | pUC18T-miniTn7T-zeo with the gene *pilA* used as a negative control for plasmid pUC18T-miniTn7T-zeo | [7] |
| **pEX18Ap** | Plasmid to generate mutants using allelic exchange and sucrose counterselection | [8] |
| **pEX18Ap_carO398** | Plasmid for generation of *carO* deletion mutants | This study |

**S8 Table.**
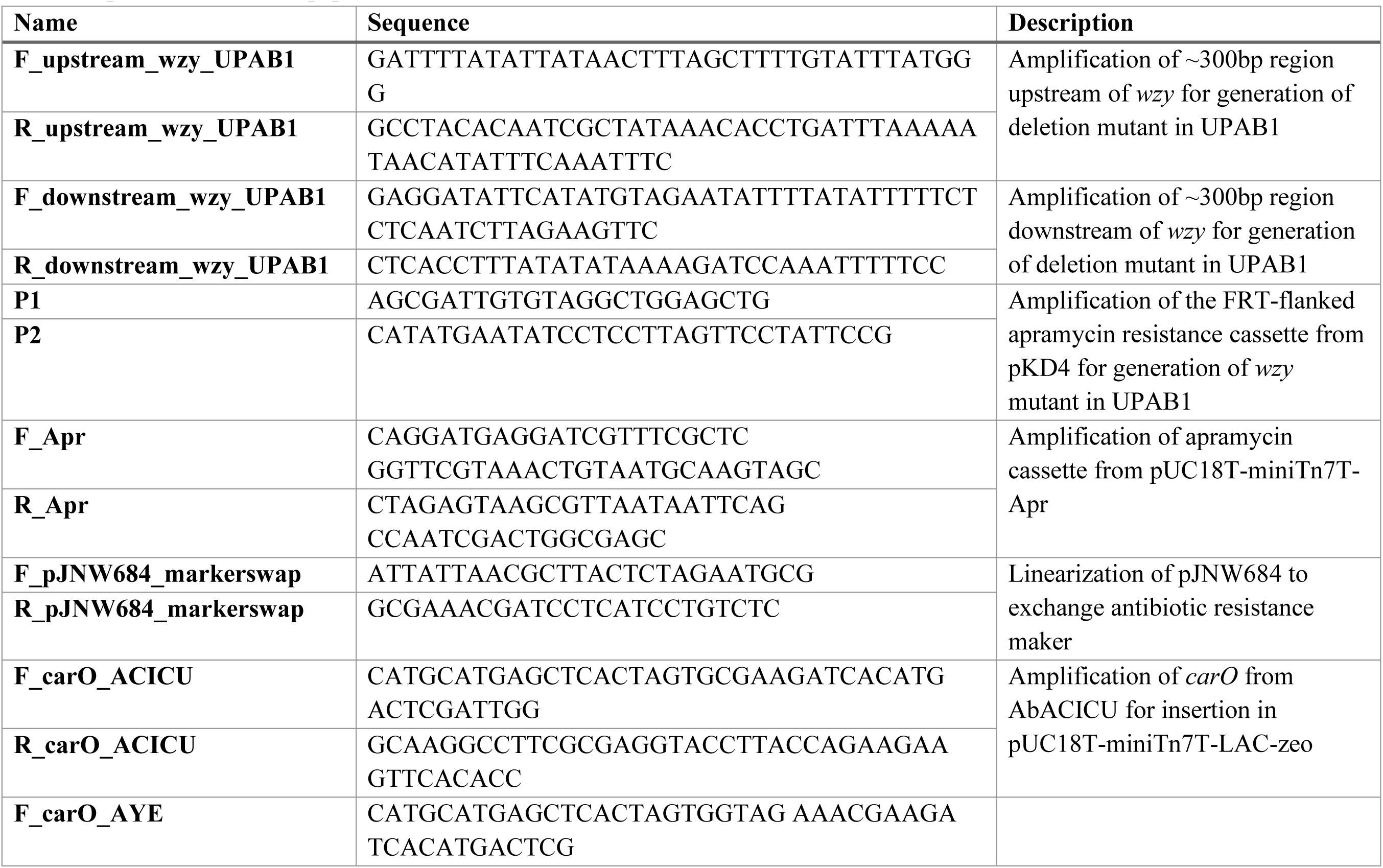
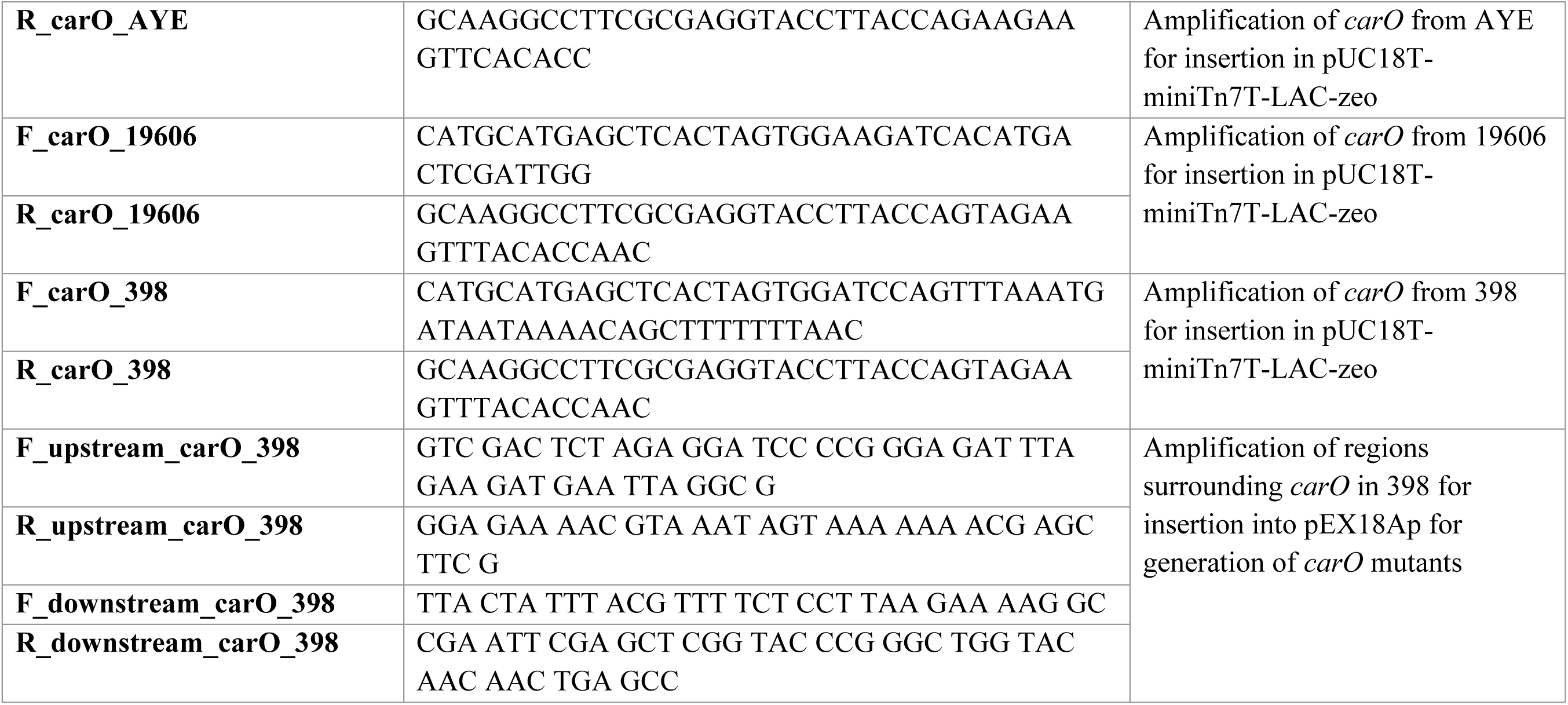
Primers. List of all primers used in the paper.

